# Involvement of PHOSPHATE TRANSPORTER TRAFFIC FACILITATOR1 in COPII assembly by interacting with SAR1 GTPase

**DOI:** 10.1101/2025.01.09.632146

**Authors:** Hui-Fang Lung, Jia-Dong Chu, Tzu-Yin Liu

**Author notes:** Address correspondence to Institute of Bioinformatics and Structural Biology, College of Life Sciences and Medicine, National Tsing Hua University, 101, Section 2, Kuang-Fu Road, Hsinchu 300044, Taiwan R.O.C., **Corresponding author:** *Tzu-Yin Liu Email:.

## Abstract

Inorganic phosphate (Pi) uptake and translocation are crucial for plant growth and development, relying on plasma membrane targeting of PHOSPHATE TRANSPORTER1 (PHT1) transporters. The plant-specific endoplasmic reticulum (ER)-resident PHOSPHATE TRANSPORTER TRAFFIC FACILITATOR1 (PHF1) is structurally related to SEC12, which initiates the coat protein complex II (COPII) assembly as a guanine nucleotide exchange factor (GEF) by activating the small GTPase SAR1. In contrast, PHF1 loses the conserved catalytic residues critical for GEF activity and specifically assists the ER exit of the PHT1 transporters. However, the underlying molecular mechanism remains unknown. In this study, we showed that overexpression of *Arabidopsis thaliana* PHT1;1 (*At*PHT1;1) in the tobacco transient expression system caused a portion of *At*PHF1 distribution into *At*SAR1b- and *At*SEC24a-labeled ER exit sites. We demonstrated that *At*PHF1 interacts with *At*SAR1b and *At*SAR1c based on the tripartite split-GFP association in agro-infiltrated tobacco leaves and verified this interaction using miniTurbo-based proximity labeling. We also confirmed its physiological relevance by co-immunoprecipitation of the endogenous *At*PHF1 with *At*SAR1c-GFP in *Arabidopsis* transgenic lines. Importantly, *At*PHF1 preferentially interacts with the GDP-locked *At*SAR1. Therefore, we propose that *At*PHF1 or the *At*PHT1;1-*At*PHF1 complex interacts with the SAR1 GTPase to participate in the early step of COPII recruitment for the ER export of PHT1 transporters.

## Introduction

As sessile organisms, plants adjust the protein and lipid composition of the plasma membrane (PM) to sense and adapt to the ever-changing environment. Such dynamic cellular processes heavily rely on the modulation of the endomembrane trafficking system (Wang et al., 2020), including the coat protein complex II (COPII)-mediated endoplasmic reticulum (ER)-to-Golgi transport pathway. Although the core COPII machinery, composed of five cytosolic factors, SAR1, SEC23, SEC24, SEC13, and SEC31, is conserved throughout all eukaryotes, the mechanisms underlying COPII recruitment and assembly are most extensively studied in the budding yeast *Saccharomyces cerevisiae* (Barlowe and Miller, 2013). In this model organism, the COPII-related components are encoded by single genes and recruited sequentially to the ER exit sites (ERES) (Kurokawa and Nakano, 2019). The SAR1 GTPase is activated upon the exchange of GDP to GTP by the guanine nucleotide exchange factor (GEF) SEC12, exposing its amphipathic N-terminal helix to be inserted into the ER membrane (d’Enfert, 1991; Paul et al., 2023). SAR1 then interacts directly with SEC23, inducing the formation of the inner coat SEC23–SEC24 complex, which in turn recruits the outer coat SEC13–SEC31 complex (Bi, 2002; Stagg et al., 2006). In addition, SEC16, an initial factor assembled at the ERES, serves as a scaffold protein by concentrating SEC12 and GTP-bound active SAR1 in proximity (Supek et al., 2002; Montegna et al., 2012), and was shown to stabilize COPII coat assembly by inhibiting SAR1 inactivation (Yorimitsu and Sato, 2012). By comparison, the plant COPII subunit paralogs are encoded by multi-gene families and outnumber those in other organisms (Chung et al., 2016). Multiple COPII paralogues allow the diversity of COPII transport carriers to adapt to various developmental and stress-related cues (Chung et al., 2016; Wang et al., 2020). For instance, the phytohormone abscisic acid triggered the formation of *Arabidopsis thaliana* SAR1a (*At*SAR1a)-dependent COPII vesicles that carry osmotic stress-related carriers (Li, 2021). The *At*SAR1a-*At*SEC23a pairing was also shown to mediate the ER export of the ER-associated transcription factor bZIP28, which is upregulated in response to ER stress (Zeng et al., 2015). By contrast, the mechanism by which the COPII machinery regulates protein ER export to cope with nutrient deficiency remains unknown.

Inorganic phosphate (Pi) is an essential macronutrient for plant growth. Due to its poor solubility and mobility in soil, Pi bioavailability is limited, thus causing plants to face Pi deficiency (Shen, 2011). Under Pi-limited conditions, plants enhance the external Pi uptake as well as the recycling and remobilization of internal Pi, which involves the upregulation of the *PHOSPHATE TRANSPORTER1* (*PHT1*) gene family at the transcript and post-transcriptional level (Nussaume et al., 2011). While most *PHT1* genes are upregulated by Pi deprivation, the newly synthesized PHT1 proteins (PHT1s) must exit the ER for the PM targeting. Similar to the inhibitory effect of SEC12 overexpression on the ER export of the Golgi transmembrane protein ERD2 in tobacco leaves (Hanton et al., 2007), the overexpression of SEC12 impaired the PM targeting of *At*PHT1;2 (Bayle et al., 2011). Thus, the ER-to-Golgi transport of PHT1s is dependent on efficient COPII assembly. Interestingly, the plant-specific SEC12-related protein PHOSPHATE TRANSPORTER TRAFFIC FACILITATOR1 (PHF1) was identified to facilitate the ER exit of PHT1s (González et al., 2005; Bayle et al., 2011). Despite sharing the sequence similarity with SEC12, *At*PHF1 lost the conserved catalytic residues critical for GEF activity and failed to rescue the growth defect of the yeast *sec12* mutant (González et al., 2005). Instead, the *Arabidopsis* loss-of-function *phf1* mutant showed decreased cellular Pi content and impaired PM targeting of *At*PHT1;1 (González et al., 2005). As the role of PHF1 in assisting the ER exit of PHT1s resembles that of the yeast Pho86p in the ER exit of Pho84p, the yeast homolog of PHT1s (Lau, 2000), PHF1 may functionally diverge from SEC12 (González et al., 2005).

The transient protein expression using tobacco leaves is a well-established plant system to study the dynamic organization of COPII proteins at the ERES (daSilva et al., 2004; Hanton et al., 2009). In this system, the ERES formation can be monitored by the subcellular distribution of fluorescent protein-tagged COPII components changing from cytosolic patterns into punctate structures (daSilva et al., 2004; Hanton et al., 2007; Hanton et al., 2008; Hanton et al., 2009). Overexpression of COPII-dependent membrane cargoes enhanced the recruitment of YFP-*At*SEC24a to the ERES as well as induced the *de novo* formation of ERES (Hanton et al., 2007). Likewise, *At*PHT1;2-CFP overexpression resulted in the recruitment of YFP-*At*SEC24a to ERES puncta, reinforcing that the ER export of *At*PHT1s is COPII-dependent (Bayle et al., 2011). Although the ER-resident *At*PHF1 failed to co-localize with *At*SEC24a, thus leading to the conclusion that *At*PHF1 is not involved in COPII recruitment (Bayle et al., 2011), it remains unclear whether PHF1 participates in cargo recognition or the selective packaging of PHT1 transporters into COPII vesicles.

Since PHF1 shares sequence similarity with SEC12 yet exhibits a distinct function, we speculated that PHF1 may engage in the COPII-dependent ER export of PHT1s on a molecular basis different from that of SEC12. To explore this hypothesis, we aimed to investigate whether *At*PHT1;1 overexpression can induce the distribution of *At*PHF1 into ERES puncta and whether *At*PHF1 can interact with COPII-related components. Using agro-infiltrated tobacco leaves, we showed that overexpression of *At*PHT1;1 triggered a portion of *At*PHF1 distribution into punctate structures that partially co-localized with *At*SAR1b and *At*SEC24a. Furthermore, based on the tripartite split-GFP association, *At*PHF1 interacted with *At*SAR1b and *At*SAR1c, but not with *At*SEC24a or other COPII-related components. Consistently, we verified the interaction of *At*PHF1 and *At*SAR1b by the proximity labeling in the transient tobacco system and demonstrated that the endogenous *At*PHF1 and *At*PHT1;1/2/3 were co-immunoprecipitated with *At*SAR1c-GFP in *Arabidopsis* transgenic lines. More importantly, we showed that, like *At*SEC12, *At*PHF1 preferentially interacted with the GDP-locked inactive form of *At*SAR1b. Our results unveiled that *At*PHF1 may act as an early regulator of COPII assembly for the ER export of *At*PHT1s through interacting with *At*SAR1 proteins (*At*SAR1s).

## Results

### Overexpression of *At*PHT1;1 triggers partial distribution of *At*PHF1 into ERES-associated punctate structures

Both the mammalian and plant SEC12 proteins are dispersed throughout the ER (Weissman et al., 2001; Bayle et al., 2011). However, the mammalian SEC12 has also been reported to be enriched at ERES (Montegna et al., 2012) and can be concentrated at the ERES by the collagen cargo receptor component cTAGE5 for the ER export of collagen (Saito et al., 2014). We thus wondered whether the SEC12-related *At*PHF1, which was reported to distribute evenly across the ER (González et al., 2005; Bayle et al., 2011), can be induced to concentrate at ERES when the ER export of *At*PHT1s is highly demanded. When expressed alone in agro-infiltrated tobacco leaves, *At*PHF1-GFP showed a reticular ER pattern, while a portion of *At*PHF1-GFP displayed punctate-like structures when the split-GFP tagged *At*PHT1;1 (*At*PHT1;1-S11) was co-expressed (**Fig. 1A**). By contrast, the co-expression of *At*PHF1-GFP with the sugar transporter *At*STP1-S11 barely changed the distribution of *At*PHF1-GFP (**Fig. 1A**). Statistical analysis further suggested that the overexpression of *At*PHT1;1-S11 but not of *At*STP1-S11 significantly triggered the partial distribution of *At*PHF1-GFP into punctate structures (**Fig. 1B**). To ensure that the formation of *At*PHF1 puncta triggered by *At*PHT1;1 overexpression is not due to the ER morphological changes, we examined the ER organization in the absence and presence of *At*PHT1;1 and *At*PHF1 overexpression. Regardless of the co-expression of *At*PHT1;1 and/or *At*PHF1, the overall morphology of the ER, labeled by the mCherry-HDEL marker, showed similar interconnected ER sheets and tubular structures throughout the cortical ER (**Supplementary Fig. S1A**). Upon the overexpression of *At*PHT1;1 and/or *At*PHF1, the ER lumen marker mCherry-HDEL-labeled puncta were occasionally identified (**Fig. 1C**). However, quantification analysis showed no significant difference in the number of mCherry-HDEL-labeled puncta, regardless of the co-expression of *At*PHT1;1 and/or *At*PHF1 (**Supplementary Fig. S1B**). These mCherry-HDEL-labeled puncta might represent ER-PM contact sites as previously reported (Kriechbaumer and Brandizzi, 2020). Alternatively, they might represent only a small percentage of mCherry-HDEL fusion proteins visualized as puncta during cycling between the ER and Golgi under steady-state conditions. To quantify these observations, we classified the *At*PHT1;1-induced puncta into three groups: the distinct *At*PHF1-labeled puncta that did not overlap with the mCherry-HDEL puncta, the distinct mCherry-HDEL-labeled puncta that did not overlap with the *At*PHF1-labeled puncta, and the overlapping puncta. Statistical analyses indicated that the distinct *At*PHF1-labeled puncta predominated over the overlapping puncta (**Fig. 1D**; detailed analysis of individual experiment in **Supplementary Fig. S2A**), suggesting that the majority of the *At*PHT1;1 overexpression-induced *At*PHF1 puncta represented a compartment different from the mCherry-HDEL-labeled puncta.

**Figure 1.**
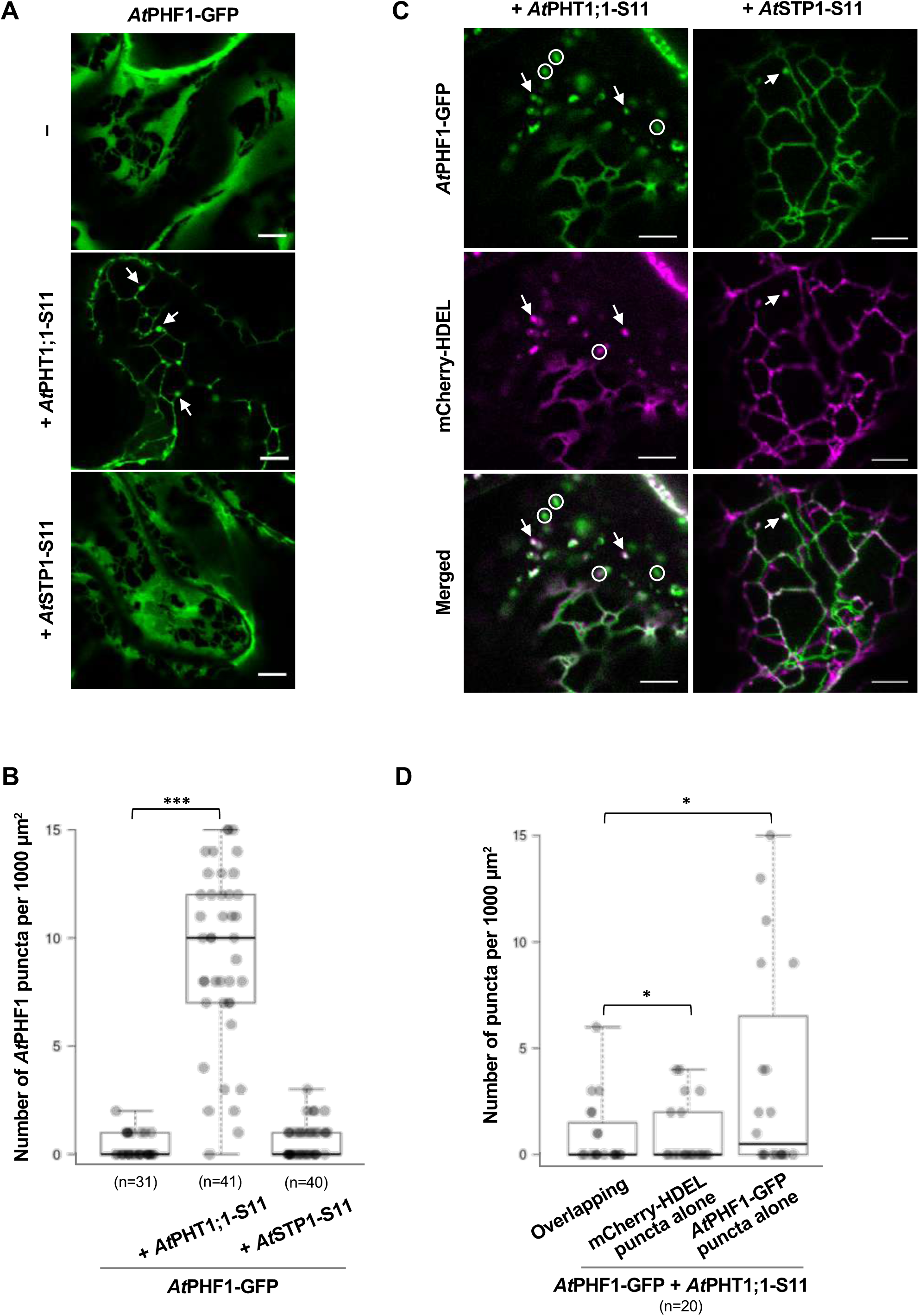
*At*PHT1;1 overexpression triggers partial distribution of *At*PHF1 to punctate structures in agro-infiltrated *N. benthamiana* leaves. (**A**) Distribution of *At*PHF1-GFP in the absence or presence of *At*PHT1;1-S11 or *At*STP1-S11 co-expression. Arrows, punctate structures. (**B**) Quantification analysis of (**A**) from three independent experiments. (**C**) Co-localization analysis of *At*PHF1-GFP puncta and mCherry-HDEL puncta in the presence of *At*PHT1;1-S11 or *At*STP1-S11 co-expression. Arrows, overlapping puncta; circles, non-overlapping puncta. (**D**) The quantification of (**C**) from three independent experiments. Representative confocal images at the peripheral layers of the epidermis are shown in (**A**) and (**C**); scale bars, 5 µm. Box plots show medians (center lines), interquartile ranges (boxes), data ranges (whiskers), and data points (dots) in (**B**) and (**D**). The number of regions of interest used for quantification is shown in parentheses. ***, *P* < 0.001 versus *At*PHF1-GFP expression alone in (**B**); *, *P* < 0.05 versus overlapping puncta in (**D**); Dunnett’s test for multiple comparisons.

In the tobacco transient expression system, the co-expression of membrane cargoes recruited the COPII components, such as *At*SAR1 and *At*SEC24, to concentrate at the punctate ERES (Hanton et al., 2007; Hanton et al., 2008). Intriguingly, the populations of *At*PHF1 puncta upon *At*PHT1;1 overexpression displayed a size with a median of 0.66–0.77 µm (**Supplementary Fig. S2B**), which is comparable to the previously reported size of ERES puncta with a range of 0.15–0.8 µm across various organisms (Bannykh, 1996; Yang et al., 2005; Zeuschner et al., 2006; Takagi et al., 2020; Weigel et al., 2021). To address whether the punctate structures of *At*PHF1-GFP induced by the overexpression of *At*PHT1;1 might represent ERES, we selected *At*SAR1b as a COPII marker to co-localize with *At*PHF1-mCherry. This is because *AtSAR1b* is the most highly expressed *AtSAR1* gene in roots and is upregulated in response to Pi deprivation (Liu et al., 2016) (**Supplementary Fig. S3**). In addition, the previous proteomic analysis of the Pi overaccumulator *pho2* root has revealed the co-induction of *At*SAR1b and *At*PHT1s (Huang et al., 2013), implying an increased demand for *At*SAR1b to promote the ER-to-Golgi traffic of *At*PHT1s. Considering that the N-terminal tail of SAR1 is inserted into the ER membranes upon activation (d’Enfert, 1991; Paul et al., 2023), we fused the S11 tag to the C-terminus of *At*SAR1b to prevent steric hindrance. In tobacco epidermal cells, *At*SAR1b-S11 exhibited both a diffuse cytosolic distribution and an ER membrane-associated punctate pattern, similar to the previously reported results (Yang et al., 2005; Hanton et al., 2008) (**Supplementary Fig. S4A**). *At*SEC24a is involved in cargo selection for COPII vesicles and is also widely used as a reliable ERES marker (Miller, 2002; Hanton et al., 2007); therefore, we generated a construct expressing *At*SEC24a-S11 for co-localization with *At*PHF1-mCherry as well. The subcellular localizations of *At*SEC24a-S11 displayed the cytosolic and punctate patterns (**Supplementary Fig. S4A**). *At*PHT1;1 overexpression induced the formation of *At*PHF1-mCherry punctate structures that co-localized with the *At*SAR1b- and *At*SEC24a-labeled ERES (**Fig. 2A** and **2B**). Statistical analyses further suggested that the majority of *At*PHF1-mCherry puncta overlapped with these two ERES markers (**Fig. 2C**; detailed analysis of individual experiment in **Supplementary Fig. S5A** and **S5B**). The size of the overlapping puncta was also comparable to that of the distinct *At*SAR1b- and *At*SEC24a-labeled puncta, with a median of 0.67–0.81 µm (**Supplementary Fig. S5C** and **S5D**). The co-localization analysis further indicated a better spatial co-occurrence of *At*PHF1-mCherry puncta with *At*SAR1b than with *At*SEC24a, as determined by a significantly higher Manders’ overlap coefficient for *At*SAR1b (**Fig. 2D**). Taken together, these results suggested that when *At*PHT1;1 is overexpressed as a COPII-dependent export membrane cargo, *At*PHF1 can partially distribute into puncta that are associated with ERES markers.

**Figure 2.**
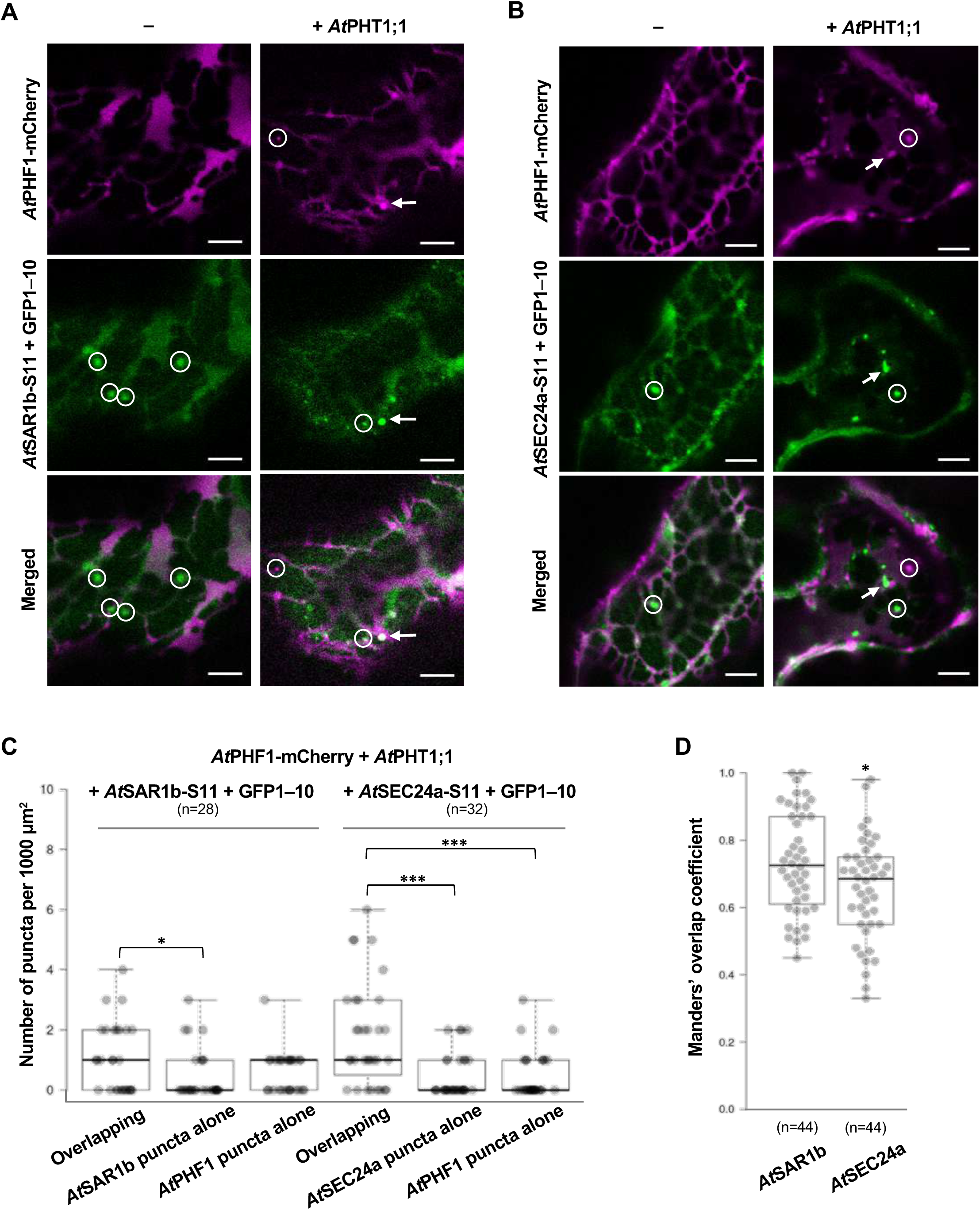
*At*PHT1;1-induced *At*PHF1 puncta partially co-localize with the ERES markers in agro-infiltrated *N. benthamiana* leaves. (**A**, **B**) Co-localization of *At*SAR1b-S11 (**A**) and *At*SEC24a-S11 (**B**) with *At*PHF1-mCherry in the absence or presence of *At*PHT1;1 overexpression. Arrows, overlapping puncta; circles, non-overlapping puncta. Scale bars, 5 µm. Representative confocal images taken at the peripheral layer of the epidermis are shown from three independent experiments. (**C**) The quantification of (**A**) and (**B**) from three independent experiments. The number of regions of interest used for quantification is shown in parentheses. *, *P* < 0.05; ***, *P* < 0.001; Dunnett’s test for multiple comparisons against overlapping with *At*SAR1b or *At*SEC24a. (**D**) Comparison of Manders’ overlap coefficient between the co-localization of *At*PHF1-mCherry with *At*SAR1b-S11 and *At*SEC24a-S11. Data were collected from three independent experiments. Box plots show medians (center lines), interquartile ranges (boxes), data ranges (whiskers), and data points (dots). The puncta numbers used for the quantification analysis are shown in parentheses. *, *P* < 0.05; two-tailed Mann-Whitney U test.

To further determine whether *At*PHF1 enhances COPII recruitment beyond the effect of *At*PHT1;1 overexpression alone, we quantified *At*SAR1b-labeled ERES puncta under (1) *At*PHF1-mCherry overexpression, (2) *At*PHT1;1 overexpression, and (3) co-expression of *At*PHF1-mCherry and *At*PHT1;1. Overexpression of *At*PHF1-mCherry alone did not induce ERES formation (**Supplementary Fig. S6A** and **S6B**), in agreement with the previous study (Bayle et al., 2011). Consistent with the notion that COPII-dependent membrane cargo can stimulate *de novo* ERES formation (Hanton et al., 2007; Bayle et al., 2011), *At*PHT1;1 overexpression increased the number of *At*SAR1b-labeled puncta (**Supplementary Fig. S6A** and **S6B**). In addition, co-expression of *At*PHT1;1 and *At*PHF1-mCherry decreased the number of *At*SAR1b-labeled ERES puncta compared to the *At*PHT1;1 overexpression alone, indicating that *At*PHF1 attenuated the *At*PHT1;1-induced increase in ERES formation (**Supplementary Fig. S6B**).

### The interaction of *At*PHF1 with *At*SAR1b/c *in planta* based on tripartite split-GFP association

As *At*PHT1;1 overexpression triggered partial distribution of *At*PHF1 into punctate structures associated with *At*SAR1b and *At*SEC24a, we postulated that *At*PHF1 may participate in COPII recruitment or assembly. To explore this possibility, we examined the interaction of *At*PHF1 with several COPII-related components based on the tripartite split-GFP association in agro-infiltrated tobacco leaves (Cabantous et al., 2013; Liu et al., 2018). The expression of split-GFP-tagged *At*SEC12 and *At*PHF1 (S10-*At*SEC12 and S10-*At*PHF1) confirmed their localization at the ER when co-expressed with the cytosolic S11-GFP1–9 (**Supplementary Fig. S4B**). We also generated constructs expressing the split-GFP-tagged *At*SAR1c, *At*SEC13a, and *At*SEC16a (*At*SAR1c-, *At*SEC13a-, and *At*SEC16a-S11). As *AtSAR1c* is the second most highly expressed *AtSAR1* gene in roots (Liu et al., 2016) (**Supplementary Fig. S3**) and plays an interchangeable role with *AtSAR1b* at the protein level in pollen development (Liang et al., 2020), we included *At*SAR1c for comparison with *At*SAR1b. As previously reported (Hanton et al., 2008; Hanton et al., 2009; Takagi et al., 2013), *At*SAR1c-S11, *At*SEC13a-S11, and *At*SEC16a-S11 predominantly localized to the cytosol and ERES puncta (**Supplementary Fig. S4A**). The expression of 3xHA-S11 or S10-3xHA exhibited cytosolic patterns (**Supplementary Fig. S4**) and was used as a negative control for the interaction (**Fig. 3A–C**). The interaction of S10-*At*SEC12 with *At*SAR1b- or *At*SAR1c-S11 was used as a positive control, yielding the GFP complementation signals (**Fig. 3A**). Notably, like S10-*At*SEC12, S10-*At*PHF1 interacted with *At*SAR1b- and *At*SAR1c-S11 but not with *At*SEC24a-, *At*SEC13a-, and *At*SEC16a-S11 (**Fig. 3B**), suggesting the specificity of *At*PHF1 interaction with *At*SAR1s among the COPII-related proteins.

**Figure 3.**
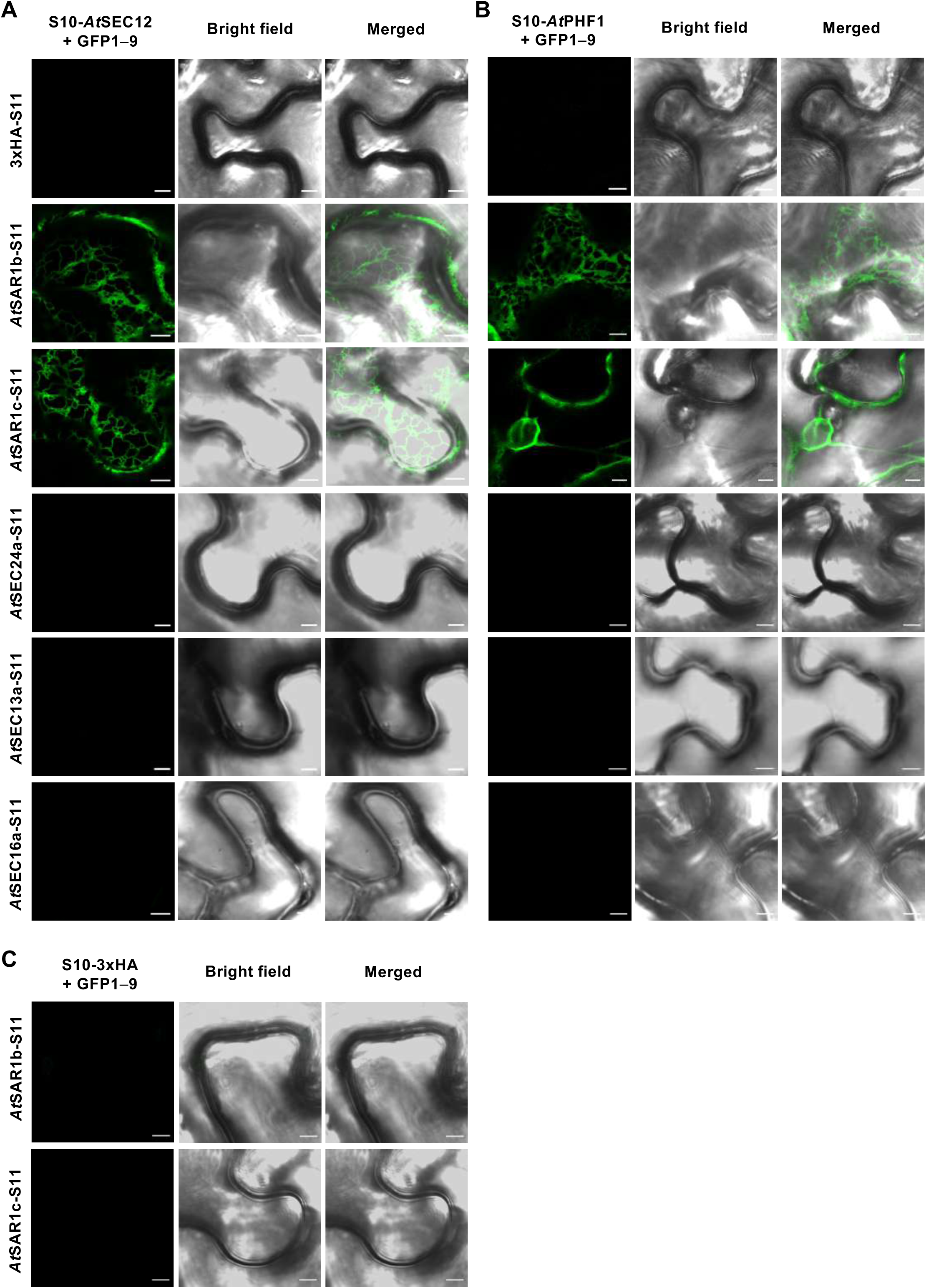
*At*PHF1 interacts with *At*SAR1b and *At*SAR1c *in planta* based on the tripartite split-GFP association in agro-infiltrated *N. benthamiana* leaves. The interaction of S10-*At*SEC12 (**A**) or S10-*At*PHF1 (**B**) with the COPII-related components, *At*SAR1b-, *At*SAR1c-, *At*SEC24a-, *At*SEC13a-, and *At*SEC16a-S11. The interactions with 3xHA-S11 in (**A**) and (**B**) and S10-3xHA (**C**) were used as negative controls. Scale bars, 5 µm. Representative confocal images taken at the peripheral layer of the epidermis are shown from three independent experiments.

### The interaction of *At*PHF1 with *At*SAR1b *in planta* based on miniTurbo (mTb) proximity labeling

The tripartite split-GFP association detects protein-protein interaction without spurious background signals even when bait and prey proteins are overexpressed (Cabantous et al., 2013; Liu et al., 2018). Still, we employed the proximity labeling method as a complementary approach to validate the *in planta* interaction between *At*PHF1 and *At*SAR1b. Proximity labeling begins after the application of biotin and can be reversibly halted by removing biotin, thereby allowing the capture of transient and dynamic protein-protein interactions at a ten-nanometer scale (Kim et al., 2014; Xu et al., 2023). Due to the superior feasibility of miniTurbo (mTb) in the tobacco system (Mair et al., 2019), the mTb-based proximity labeling was used. For unknown reasons, our initial attempt to express mTb-EYFP-*At*PHF1 in agro-infiltrated tobacco leaves was unsuccessful, so we generated the *At*SAR1b-mTb-S11 construct by fusing mTb to *At*SAR1b (**Fig. 4A**). We used the cytosol-localized mTb-NES-EYFP (**Fig. 4B**), which carries the nuclear export sequence (NES), as a negative control (Mair et al., 2019). Taking advantage that S11 can self-complement with GFP1–10, we could detect the expression and subcellular distribution of *At*SAR1b-mTb-S11 in agro-infiltrated tobacco leaves (**Fig. 4B**). We then co-expressed mTb-NES-EYFP or *At*SAR1b-mTb-S11 with *At*SEC12-mCherry or *At*PHF1-mCherry and incubated the harvested leaves in a biotin solution, followed by tissue homogenization and protein extraction. Protein extraction from non-infiltrated tobacco leaves was used for the background comparison. As the expression of the cytosolic mTb-NES-EYFP was much greater than the membrane-associated *At*SAR1b-mTb-S11, we adjusted the ratio of protein amount for *At*SAR1b-mTb-S11: mTb-NES-EYFP to 30: 1 (**Supplementary Fig. S7**). In this way, the protein amounts of *At*SAR1b-mTb-S11 and mTb-NES-EYFP were comparable, allowing them to compete fairly for binding to streptavidin beads. When co-expressed with *At*SAR1b-mTb-S11, both *At*SEC12-mCherry and *At*PHF1-mCherry could be captured by streptavidin and detected by immunoblot, indicating that they were biotinylated (**Supplementary Fig. S7**). By contrast, the co-expression of mTb-NES-EYFP with *At*SEC12-mCherry or *At*PHF1-mCherry did not yield similar effects (**Supplementary Fig. S7**). Differences in protein loading ratios may artificially enhance or mask interactions. Therefore, we also used equal amounts of protein in the experiments. Consistently, even though the expression of *At*SAR1b-mTb-S11 was over five times lower than that of mTb-NES-EYFP, about two times more *At*SEC12-mCherry or *At*PHF1-mCherry was captured by streptavidin when co-expressed with *At*SAR1b-mTb-S11 (**Fig. 4C**). These results again suggested that *At*PHF1 interacts with *At*SAR1s *in planta*.

**Figure 4.**
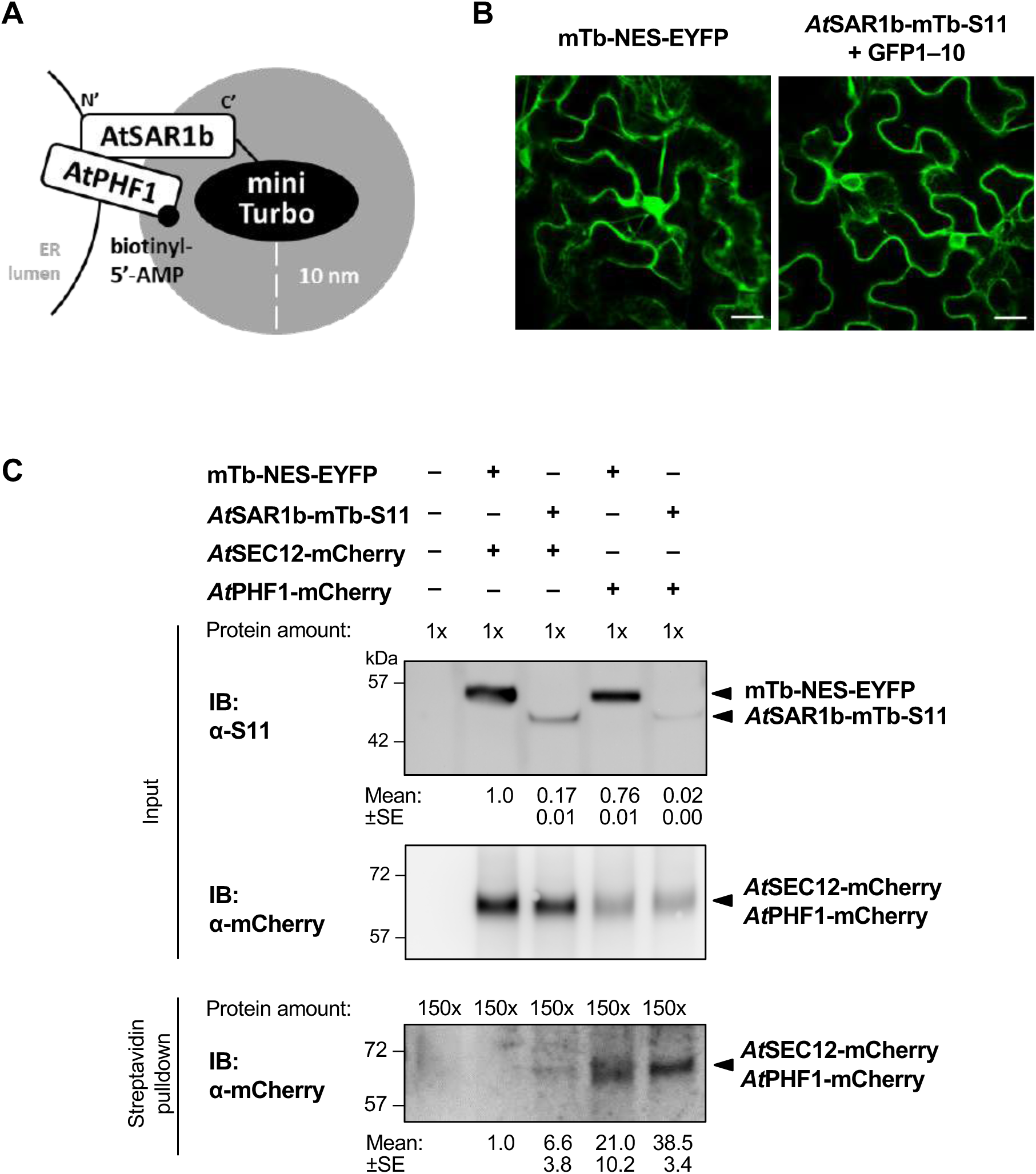
*At*PHF1 interacts with *At*SAR1b *in planta* based on the proximity labeling in agro-infiltrated *N. benthamiana* leaves. **(A)** Schematic diagrams of mTb-fused bait protein in the proximity of the prey protein. The reactive intermediate biotinyl-5’-AMP biotinylates proximal prey proteins within the indicated range (grey zone). **(B)** The expression and subcellular distribution of mTb-NES-EYFP and *At*SAR1b-mTb-S11. Scale bars, 20 µm. Representative confocal images are shown from three independent experiments. **(C)** The interaction analysis of *At*PHF1-mCherry and *At*SAR1b by proximity labeling. Equal amounts of protein from samples were used for streptavidin pulldown. Anti-S11 antibody was used to detect EYFP-tagged and S11-tagged fusion proteins. The relative intensity is shown as mean ± SE from two independent experiments.

To determine which region of *At*PHF1 interacts with *At*SAR1s, we generated four N-terminally S10-tagged *At*PHF1 truncated variants as follows: the cytoplasmic domain alone (*At*PHF1 N), the cytoplasmic domain with the transmembrane (TM) domain (*At*PHF1 N–TM), the TM domain alone (*At*PHF1 TM), and the TM domain with the ER-luminal domain (*At*PHF1 TM–C) (**Supplementary Fig. S8A**). The expression and subcellular distribution of these *At*PHF1 variants suggested that the S10-*At*PHF1 N variant was present in the cytosol and nucleus, while the other truncated forms of *At*PHF1 localized to the ER (**Supplementary Fig. S8B**). The subcellular localization of the C-terminally S11-tagged truncated forms of *At*PHF1 also showed similar results (**Supplementary Fig. S8C**), suggesting that the TM domain of *At*PHF1 alone is sufficient for its ER targeting. Of note, the interaction of S10-tagged *At*PHF1 N or *At*PHF1 N–TM with *At*SAR1b- or *At*SAR1c-S11 displayed both the ER and puncta patterns (**Supplementary Fig. S9A** and **S9B**). By contrast, the interaction of S10-tagged *At*PHF1 TM or *At*PHF1 TM–C with *At*SAR1b- or *At*SAR1c-S11 additionally resulted in a few GFP signals that did not display a typical reticular ER pattern (**Supplementary Fig. S9A** and **S9B**). These observations suggest that both the cytosolic and the transmembrane domains of *At*PHF1 contribute to the interaction with *At*SAR1s at the ER.

### Co-immunoprecipitation of endogenous *A*tPHF1 with *At*SAR1c-GFP in *Arabidopsis* root

To determine whether the interaction of *At*PHF1 and *At*SAR1s is physiologically relevant, we used *Arabidopsis* transgenic plants expressing *At*SAR1c-GFP or GFP under the UBQ10 promoter for co-IP analysis (Zeng et al., 2015). The *Arabidopsis* seedlings were subjected to seven days of Pi starvation to mimic the physiological conditions in which endogenous *At*PHF1 is upregulated and the ER export of *At*PHT1s is also highly demanded. A total root protein extract was used for GFP-Trap co-IP, followed by immunoblot analysis using an anti-*At*PHF1 antibody. Due to the differences in the expression level between the cytosolic GFP and the membrane-associated *At*SAR1c-GFP in transgenic plants, we increased the protein amount for the co-IP of *At*SAR1c-GFP by 50-fold compared to that for the co-IP of GFP. Immunoblot results indicated that endogenous *At*PHF1 was co-immunoprecipitated with *At*SAR1c-GFP but not with GFP (**Supplementary Fig. S10**). As mentioned above, we also used equal amounts of root total proteins for co-IP analysis to minimize potential artifacts. The expression of *At*SAR1c-GFP was ten times lower than that of GFP; however, *At*SAR1c-GFP co-immunoprecipitated about three times more endogenous *At*PHF1 compared to GFP (**Fig. 5A**), suggesting that *At*SAR1c-GFP exhibits a higher binding affinity toward *At*PHF1. Moreover, endogenous *At*PHT1;1/2/3 proteins could also be co-immunoprecipitated with *At*SAR1c-GFP (**Fig. 5A** and **Supplementary Fig. S10**), indicating the presence of a protein complex containing *At*SAR1s, *At*PHF1, and *At*PHT1s. To dissect whether the interaction of *At*PHF1 and *At*SAR1 is direct or indirect, we performed the yeast membrane-based split-ubiquitin (mbSUS) assay. Consistent with previous reports (Huang et al., 2013; Chiu et al., 2026), *At*PHT1;1 interacted with both *At*PHF1 and the *Arabidopsis* CORNICHON HOMOLOG 5 (*At*CNIH5), a recently identified ER cargo receptor of *At*PHT1s (**Fig 5B**). *At*PHT1;1 also interacted with *At*SAR1c but not with the PM-localized organic cation/carnitine transporter *At*OCT1 (**Fig 5B**), indicating that *At*PHT1;1 interacts directly with *At*SAR1. For unknown reasons, we failed to detect the interaction of *At*PHF1 or *At*SEC12 with *At*SAR1s in the yeast mbSUS (**Supplementary Fig. S11A** and **S11B**).

**Figure 5.**
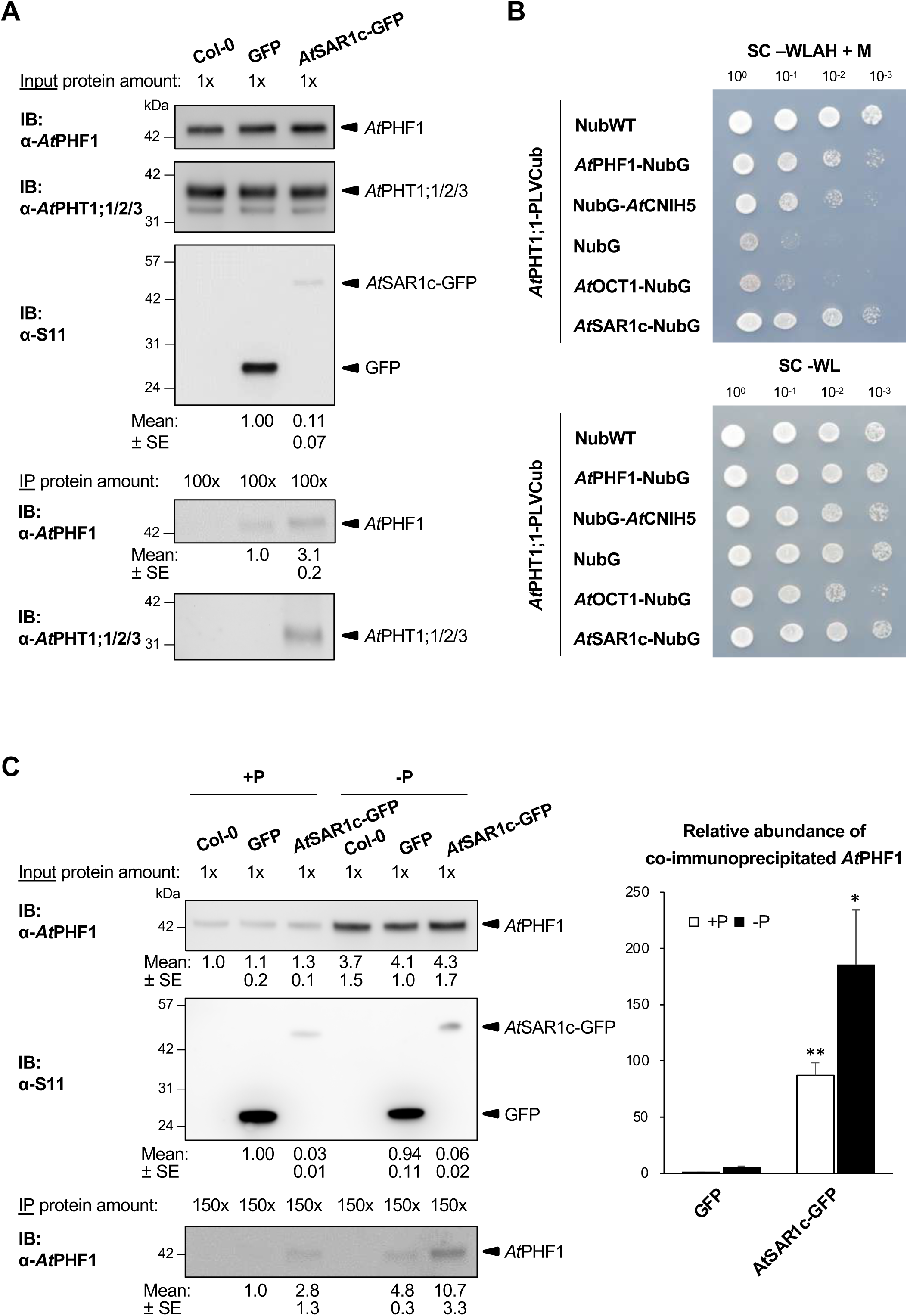
Co-immunoprecipitation of the endogenous *At*PHF1 and *At*PHT1;1/2/3 with *At*SAR1c-GFP in *Arabidopsis* transgenic plants and interaction analysis of AtSAR1c -AtPHT1;1 in the yeast split-ubiquitin system. **(A)** The interaction analyses of *At*SAR1c-GFP with the endogenous *At*PHF1 and *At*PHT1;1/2/3 in 11-day-old wild-type (Col-0), *UBQ10:*GFP, and *UBQ10:At*SAR1c-GFP seedlings subjected to Pi deprivation (0 µM KH_2_PO_4_, 7 days of starvation). **(B)** Interaction analysis of *At*SAR1c with *At*PHT1;1 in the yeast split-ubiquitin system. Co-expressions of *At*PHT1;1-CubPLV with NubWT, *At*PHF1-NubG, and NubG-*At*CNIH5 were used as positive controls, while the co-expressions with NubG and *At*OCT1-NubG were used as negative controls. Yeast transformants were grown on synthetic medium lacking tryptophan and leucine (SC–WL) for growth detection or on synthetic medium lacking tryptophan, leucine, adenine, and histidine containing 0.5 µM methionine (SC–WLAH + M) for testing protein-protein interaction. Representative results are shown from two independent experiments. **(C)** The interaction analysis of *At*SAR1c-GFP with endogenous *At*PHF1 in 11-day-old Col-0, *UBQ10:*GFP, and *UBQ10:At*SAR1c-GFP seedlings under Pi-sufficient (+P) and Pi-deficient (–P; 0 µM KH_2_PO_4_, 7 days of starvation) conditions. The relative abundance of co-immunoprecipitated endogenous *At*PHF1 proteins was calculated by normalizing them with the corresponding GFP control under +P or –P conditions. Error bars represent standard error (SE) (n = 3, independent experiments). Data significantly different from corresponding *UBQ10:*GFP are indicated by asterisks (*, *P* < 0.05; **, *P* < 0.05; two-tailed Student’s t-test). Equal amounts of protein from samples were used for GFP-Trap co-immunoprecipitation. Anti-S11 antibody was used to detect GFP fusion proteins and the relative intensity is shown as mean ± SE from three independent experiments in **(A)** and **(C)**.

It is well established that inositol pyrophosphates (PP-InsPs) respond to the varying levels of Pi in tissues. These changes modulate the interaction between SPX domains and PHR transcription factors or P1BS DNA, thereby regulating phosphate homeostasis in plants (Ried et al., 2021; Whitfield et al., 2026). To determine if a similar situation applies to the interaction between *At*PHF1 and *At*SAR1, we conducted co-IP assays in transgenic plants grown under Pi sufficiency and deficiency. Regardless of Pi status, *At*SAR1c-GFP co-immunoprecipitated endogenous *At*PHF1, as determined by normalization to the corresponding GFP control (**Fig. 5C**). The co-immunoprecipitation of *At*PHF1 with *At*SAR1c-GFP was nearly 30-fold higher compared to that with GFP during Pi-deficiency (**Fig. 5C**), consistent with our previous results (**Fig. 5A**). Similarly, *At*SAR1c-GFP also co-immunoprecipitated *At*PHF1 under Pi sufficiency (**Fig. 5C**), supporting an interaction between *At*PHF1 and *At*SAR1c under different Pi regimes. Both GFP and *At*SAR1c-GFP showed increased co-immunoprecipitation of *At*PHF1 during Pi deficiency compared to Pi sufficiency. Specifically, during Pi deficiency, GFP co-immunoprecipitated *At*PHF1 with a 5-fold increase, while *At*SAR1c-GFP showed a 2-fold increase (**Fig. 5C**). As the protein expression of *At*PHF1 and *At*SAR1c-GFP increased during Pi deficiency compared to Pi sufficiency, the rise in co-immunoprecipitated *At*PHF1 during Pi deficiency may be attributed to the modest upregulation of *At*PHF1 and *At*SAR1c-GFP under these conditions. Therefore, our co-IP results suggested that the interaction between *At*PHF1 and *At*SAR1 is not modulated by Pi status.

### *At*PHF1 preferentially interacts with the GDP-locked inactive form of *At*SAR1b

Because the assembly and disassembly of COPII is controlled by the SAR1 GTPase cycle (Van der Verren and Zanetti, 2023), we next asked whether *At*PHF1 interacts with *At*SAR1s in a specific nucleotide-binding state. Confocal analysis suggested that both the wild-type *At*SAR1b-S11 and the GTP-locked *At*SAR1b[H74L]-S11 localized to the ER membrane and cytosol as previously reported (daSilva et al., 2004; Wei and Wang, 2008) (**Supplementary Fig. S4A** and **S12A**). The GDP-locked *At*SAR1b[T34N]-S11 also displayed a similar subcellular pattern (**Supplementary Fig. S12A**). Furthermore, to quantify the soluble and membrane-associated populations of the *At*SAR1b variants, we applied biochemical fractionation. Although the microsomal fraction was slightly contaminated with the soluble fraction, as indicated by the detection of a small amount of cytosolic FBPase, the majority of the *At*SAR1b proteins were membrane-associated (**Supplementary Fig. S12B**). Namely, the membrane-associated proportions of *At*SAR1b-S11, *At*SAR1b[H74L]-S11, and *At*SAR1b[T34N]-S11 were 76.4%, 96.1%, and 67.7%, respectively, suggesting that the GDP-locked form of *At*SAR1b is relatively less membrane-associated (**Supplementary Fig. S12B**). These results were consistent with the previous finding that upon GTP binding, SAR1 is activated to insert into the ER membrane (d’Enfert, 1991; Paul et al., 2023). More importantly, the *in-planta* tripartite split-GFP assay suggested that, like *At*SEC12, *At*PHF1 preferentially interacts with the GDP-locked form of *At*SAR1b (**Fig. 6A** and **6B**). The interaction of *At*PHF1 N with the *At*SAR1b or the GDP-locked *At*SAR1b[T34N] showed cytoplasmic, ER, and puncta patterns (**Fig. 6B**). The interaction of *At*PHF1 N–TM with WT or GDP-locked *At*SAR1b was mainly localized to the ER and puncta (**Fig. 6B**). By contrast, the interaction of *At*PHF1 TM or *At*PHF1 TM–C variants with WT or GDP-locked *At*SAR1b exhibited reticular ER and punctate patterns, along with additional localization signals that were not confined to the ER (**Fig. 6B**). Overall, the GTP-locked *At*SAR1b[H74L] drastically reduced the interaction with the full-length *At*PHF1, *At*PHF1 N and *At*PHF1 N–TM variants (**Fig. 6B**). Compared to the *At*SAR1b, the GTP-locked *At*SAR1b[H74L] slightly increased the interaction with the *At*PHF1 TM and the *At*PHF1 TM–C, which was likely due to a better membrane association of *At*SAR1b[H74L] (**Fig. 6B** and **Supplementary Fig. S12B**). We interpret these as indicating that the removal of the N-terminal cytosolic domain of *At*PHF1 allows an interaction of *At*PHF1’s transmembrane domain with the GTP-locked *At*SAR1b[H74L]. We suspected that the interaction between the transmembrane domain of *At*PHF1 and the membrane-associated *At*SAR1b might occur through amino acid sequence-based recognition after *At*SAR1b is inserted into the membrane. Alternatively, this might arise from non-sequence-specific interactions in the ER membrane, potentially due to overexpression of membrane proteins. Taken together, these results indicate that the *At*PHF1 variants containing the cytosolic domain preferentially associate with the GDP-bound *At*SAR1 and thus likely participate in recruiting COPII components to ERES. To further support this finding, we performed co-IP assays in transgenic plants expressing *dex:At*SAR1b[H74L]-CFP (Lupanga et al., 2020). Upon dexamethasone (DEX) induction, *At*SAR1b[H74L]-CFP was expressed and detected in the root total protein extracts (**Fig. 6C**). When equal amounts of expressed *At*SAR1c-GFP and GTP-locked *At*SAR1b[H74L]-CFP proteins were used for IP analysis, four times more endogenous *At*PHF1 was co-immunoprecipitated with *At*SAR1c-GFP than with *At*SAR1b[H74L]-CFP (**Fig. 6C**). These results suggest that GTP-locked *At*SAR1b[H74L]-CFP has a lower binding affinity than *At*SAR1c-GFP toward *At*PHF1 in *Arabidopsis* roots.

**Figure 6.**
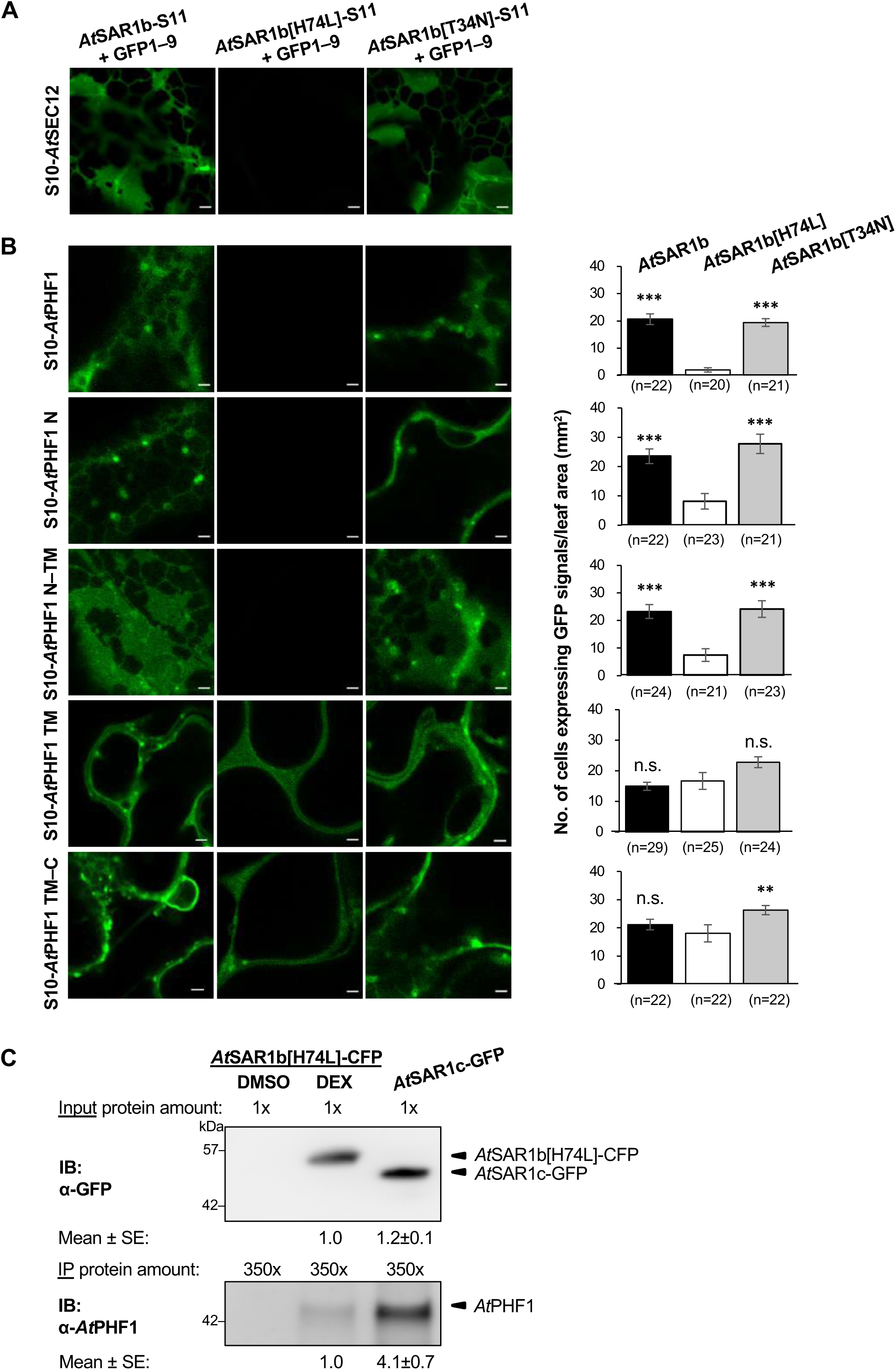
*At*PHF1 preferentially interacts with the GDP-locked form of *At*SAR1 in agro-infiltrated *N. benthamiana* leaves and *Arabidopsis* transgenic plants. (**A**, **B**) The interaction analyses of S10-*At*SEC12 (**A**) and S10-*At*PHF1 variants (**B**) with *At*SAR1b/[H74L]/[T34N]-S11 in agro-infiltrated tobacco leaves. Representative confocal images taken at the peripheral layer of the epidermis are shown from three independent experiments. Scale bars, 2 µm. Quantification indicates the number of cells expressing GFP signals per unit leaf area (mm^2^). Data were collected from three independent experiments; the number of confocal images (0.32 x 0.32 mm^2^) used for quantification is shown in parentheses. **, *P* < 0.01; ***, *P* < 0.001; not significant, n.s.; Dunnett’s test for multiple comparisons against *At*SAR1b[H74L]. (**C**) Interaction analyses of the WT or GTP-locked form of *At*SAR1 with the endogenous *At*PHF1 were performed in 11-day-old *UBQ10:At*SAR1c-GFP and *dex:At*SAR1b[H74L]-CFP seedlings subjected to Pi deprivation (0 µM KH₂PO₄, 7 days). *dex:At*SAR1b[H74L]-CFP seedlings were treated with 24 hours of 10 µM dexamethasone (DEX) or DMSO. The protein amounts as input for GFP-Trap co-immunoprecipitation are as indicated above the image. Anti-GFP antibody was used to detect GFP and CFP fusion proteins. The relative intensity is shown as mean ± standard error (SE) from two independent experiments.

## Discussion

Although several ER accessory proteins have been identified as participating in the ER exit of membrane proteins, the underlying molecular mechanisms are largely unclear (Tang et al., 2005; Bisnett et al., 2021). In *Arabidopsis*, AUXIN RESISTANCE4 (AXR4) was shown to interact with auxin influx carriers AUX1 and LAX2 to promote their PM targeting (Dharmasiri, 2006; Tidy et al., 2024). The KAONASHI3 (KNS3) family was recently identified to mediate the ER exit of the boric acid channel *At*NIP5;1 (Zhang et al., 2024). While *At*PHF1 is required for the PM targeting of *At*PHT1s, it was proposed to act as an ER-localized chaperone for the protein stability of *At*PHT1s (González et al., 2005) and not participate in the COPII vesicle formation (Bayle et al., 2011). On the contrary, our data revealed a role for *At*PHF1 in the early step of COPII assembly through the interaction with *At*SAR1s. Using three different methods—tripartite split-GFP complementation, proximity labeling, and co-immunoprecipitation—we demonstrated that *At*PHF1 interacts with *At*SAR1s *in planta* (**Fig. 3**, **4**, and **5**). Furthermore, the tripartite split-GFP and the co-IP assays demonstrated that *At*PHF1 preferentially interacts with the GDP-bound *At*SAR1s (**Fig. 6**). Interestingly, the yeast mbSUS system showed that *At*PHT1;1 could directly interact with *At*SAR1 (**Fig. 5B**). The recruitment of COPII to ERES is increased in response to the demand for ER export of membrane cargoes (Hanton et al., 2007). Our results showed that the transient overexpression of *At*PHT1;1 in *N. benthamiana* leaves also triggered partial distribution of *At*PHF1 into the punctate structures associated with the ERES markers *At*SAR1b and *At*SEC24a (**Fig. 1** and **2**). In other words, under an increased demand for the COPII-mediated export of *At*PHT1s, *At*PHF1 may partially relocate to ERES to promote this process. Although the majority of *At*PHT1;1-induced *At*PHF1 puncta co-localized with ERES markers, a small population of *At*PHF1 puncta did not co-localize with *At*SAR1b or *At*SEC24a-labeled ERES (**Fig. 2C**). These puncta were unlikely to be caused by altered ER organization, as the overall ER morphology was not altered upon co-expression of *At*PHT1;1 and *At*PHF1 (**Supplementary Fig. S1**). These puncta might arise from localized zones of high membrane protein density resulting from the co-overexpression of *At*PHT1;1 and *At*PHF1, a phenomenon known as the crowding effect (Zhou, 2009). They could also represent the dynamic ER subdomain residing by the *At*PHT1;1-*At*PHF1 complex prior to *At*SAR1 recruitment. Alternatively, they could consist of a population of *At*PHF1 that remained nearby even after the *At*PHT1-containing COPII vesicle budded away from the ERES and the ERES disappears. This possibility aligns with the role of *At*PHF1 in facilitating ER export of *At*PHT1;1, thereby reducing cargo accumulation within ERES and consequently diminishing cargo-dependent *de novo* formation of ERES (**Supplementary Fig. S6B**).

The yeast ER membrane-localized chaperone Shr3 is specifically required for proper folding of amino acid permeases and their targeting to the plasma membrane (Kota and Ljungdahl, 2005). Interestingly, the amino acid permease Gap1 split into N- and C-terminal portions can be assembled to form a functional permease in an Shr3-dependent manner (Kota et al., 2007). We also generated the N- and C-terminal portions of *At*PHT1;1 (the *At*PHT1;1 TM1–6 half and the *At*PHT1;1 TM7–12 half) to test whether *At*PHF1 plays a critical role in enabling *At*PHT1;1 to fold and attain a proper structure competent for ER export. Even upon overexpression of *At*PHF1, both the *At*PHT1;1 N and C-halves were unable to exit the ER (**Supplementary Fig. S13A**). In this scenario, neither the N-half nor the C-half of *At*PHT1;1 resulted in a partial distribution of *At*PHF1 to punctate structures (**Supplementary Fig. S13B** and **S13C**). This indicates that partial recognition of *At*PHT1;1 by *At*PHF1 is insufficient; instead, *At*PHT1;1 must attain a functional conformation that enables it to interact properly with *At*PHF1, thereby triggering a signal for ER export.

Although *At*PHF1 lacks the conserved residues required for GEF activity and cannot complement the yeast *sec12* growth defect (González et al., 2005), we surmised that upon *At*PHT1;1 overexpression, the distribution of *At*PHF1 to the ERES may facilitate the COPII recruitment or assembly. We reasoned that without co-expression of *At*PHT1s, observing partial co-localization of *At*PHF1 with ERES is beyond the detection limit of confocal imaging, which may explain the discrepancy between our results and those from a previous study (Bayle et al., 2011). In addition, *At*PHF1 interacted with *At*SAR1b and *At*SAR1c but not with other COPII-related components (**Fig. 3B**), and preferentially interacted with the GDP-locked form of *At*SAR1 *in planta* (**Fig. 6**), indicating that *At*PHF1 participates in the early step of COPII assembly, thus likely indirectly assisting the loading of *At*PHT1s into COPII vesicles for the ER export. In support of this notion, we demonstrated that endogenous *At*PHF1 and *At*PHT1;1/2/3 proteins were co-immunoprecipitated with *At*SAR1c-GFP (**Fig. 5A** and **C**), suggesting the physiological relevance of the interaction between *At*PHF1 and *At*SAR1s.

Moreover, we detected the interaction between *At*PHT1;1 and *At*SAR1c, rather than between *At*PHF1 and *At*SAR1c, using the yeast mbSUS system (**Fig. 5B**). In addition, we failed to detect the interaction of *At*SAR1s with *At*SEC12 or *At*PHF1 in the yeast mbSUS system (**Supplementary Fig. S11**), nor the interaction between the N-terminal cytosolic region of *At*PHF1 and the GDP- or GTP-bound forms of *At*SAR1b in the yeast two-hybrid (**Supplementary Fig. S14**). These negative results may reflect limitations of the heterologous system. It is also possible that the *At*PHT1;1-*At*SAR1 interaction may be more stable or direct than the *At*PHF1-*At*SAR1 interaction, which could be transient or dependent on a native ER membrane environment. The direct interaction between SAR1 and its cargoes has been reported. Mammalian SAR1 directly interacted with the di-acidic export signal of vesicular stomatitis virus glycoprotein (VSV-G) and the di-basic motif of glycolipid glycosyltransferases (GGT) (Aridor, 1998, 2001; Giraudo and Maccioni, 2003; Quintero et al., 2010). Such cargo-SAR1 interactions enhanced SEC23/24 recruitment, thereby promoting COPII coat assembly and cargo export. It remains to investigate whether the di-acidic motif within the cytoplasmic tail of *At*PHT1s interacts with *At*SAR1 for efficient ER export.

Taken together, our findings indicated that *At*PHF1 could link the specific recognition of *At*PHT1s with the modulation of *At*SAR1s activity for the COPII assembly. Such a mechanism differs from that reported for SED4, the yeast structural homolog of SEC12, which also lacks GEF activity (Gimeno, 1995; Kodera et al., 2011). Genetic and biochemical evidence suggested that SED4 and SEC12 are not functionally interchangeable and that SED4 weakly inhibits the GTPase-activating (GAP) activity of SEC23 toward SAR1 (Saito-Nakano and Nakano, 2000). On the contrary, an *in vitro* study showed that SED4 bound more strongly to the non-hydrolyzable GTP analog GMPPNP-bound SAR1 and stimulated SEC23 GAP activity, thereby accelerating the dissociation of COPII coats from the ER membrane (Gimeno, 1995; Kodera et al., 2011). While the interaction between SED4 and SEC16 is required for the localization of SEC16 to ERES (Gimeno, 1995), the interaction between *At*PHF1 and *At*SEC16a was not observed (**Fig. 3B**). Therefore, we propose that *At*PHF1 likely mediates the assembly of COPII vesicles through interacting with *At*SAR1s, thereby facilitating the ER export of *At*PHT1s. It is also likely that *At*PHF1 enhances the incorporation of *At*PHT1s into COPII vesicles by assisting the efficient formation of the *At*SAR1s-*At*PHF1-*At*PHT1s complex that participates in the ER export of *At*PHT1 (**Fig. 7**). Nevertheless, we do not exclude the possibility that PHF1 plays dual roles, acting as a membrane chaperone for the correct folding of newly synthesized PHT1s in the ER and also mediating COPII assembly. Without uncoupling these two potential functions of PHF1, its role as a chaperone at an earlier step will mask the involvement of PHF1 in the re-localization of PHT1s to ERES at a later step.

**Figure 7.**
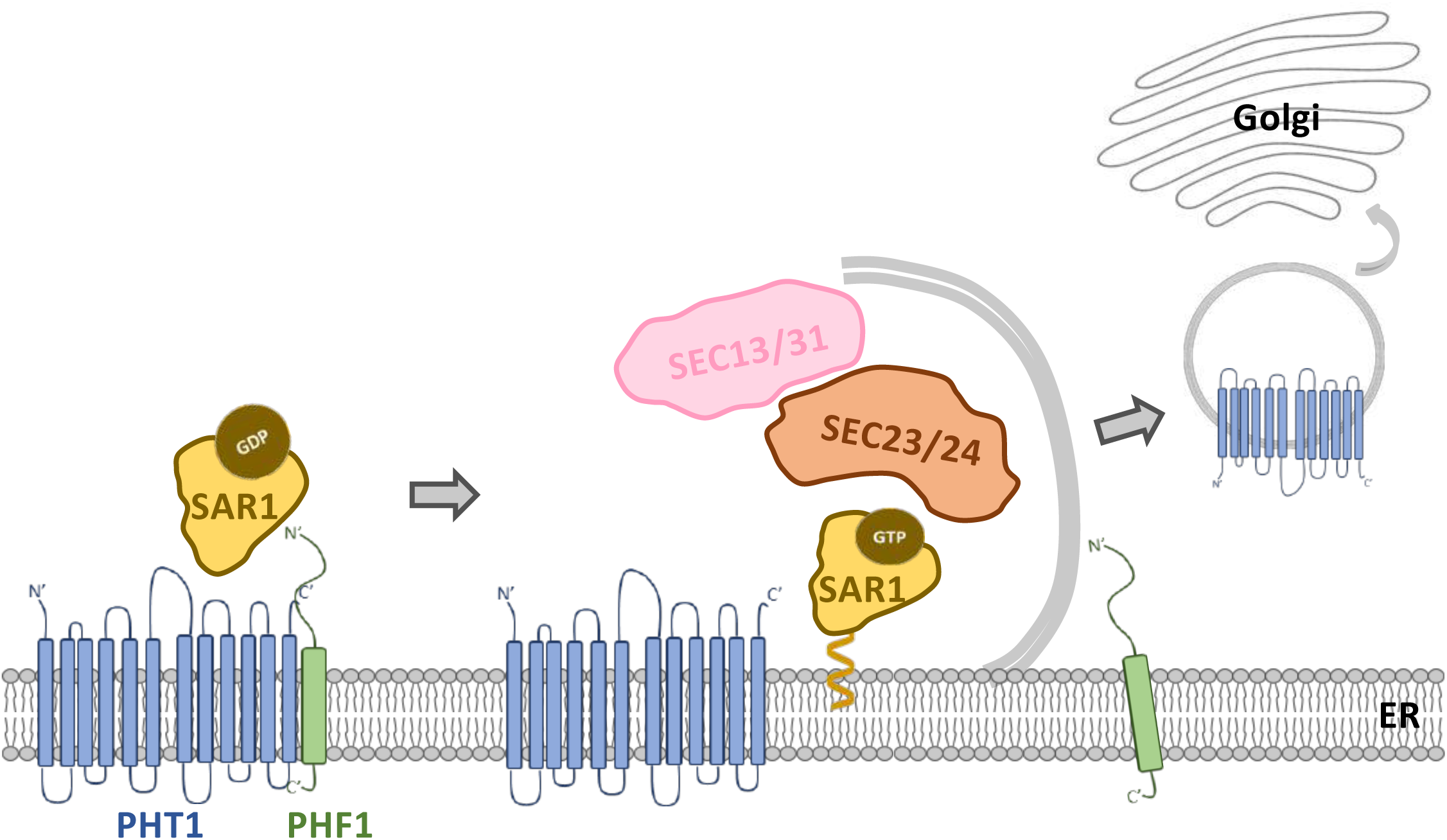
Hypothetical model depicting a role for *At*PHF1 in the early step of COPII assembly for the ER export of *At*PHT1 transporters. PHF1, or the PHT1-PHF1 complex, induces the recruitment of SAR1 to the ER exit sites (ERES), where PHF1 preferentially interacts with GDP-bound SAR1. The PHF1-SAR1 interaction may facilitate COPII assembly or stabilize the transient SAR1-PHT1-PHF1 complexes. Following activation by the guanine nucleotide exchange factor SEC12, the GTP-bound SAR1 recruits the SEC23/24 inner coat and the outer coat SEC13/31. After several rounds of SAR1 GTP hydrolysis, PHT1-containing COPII vesicles bud from the ER, and PHF1 may remain in the vicinity.

Interestingly, we recently reported that *At*CNIH5 acts as an ER cargo receptor that cycles between the ER and the early Golgi and selects *At*PHT1s for ER export (Liu et al., 2025; Chiu et al., 2026). As the *cnih5 phf1* mutant shows more severe growth defects than the *phf1* single mutant, and *At*PHF1 proteins are increased in the root of *cnih5* (Chiu et al., 2026), we suspected that *At*PHF1 and *At*CNIH5 work interdependently. Namely, while *At*PHF1 couples the recognition of *At*PHT1s and the recruitment of COPII proteins by interacting with *At*PHT1s and *At*SAR1s, *At*CNIH5 is involved in the selective packaging of *At*PHT1 into COPII vesicles.

*At*PHF1 is structurally related to SEC12 proteins, which adopt a seven-bladed β propeller (González et al., 2005; Joiner and Fromme, 2021). Based on the crystal structure, the catalytically critical K loop within the cytoplasmic domain of yeast SEC12 makes direct contact with nucleotide-free SAR1 (McMahon et al., 2012; Joiner and Fromme, 2021). By contrast, our data showed that both the cytosolic and the transmembrane domains of *At*PHF1 interact with *At*SAR1b/c *in planta* (**Supplementary Fig. S9A** and **S9B**), indicating that the interaction interface may span relatively large areas. Since the catalytic residues required for GEF activity are not conserved in *At*PHF1 (González et al., 2005), it remains to be elucidated whether the interaction interface of PHF1-SAR1s is different from that of SEC12-SAR1s and how PHF1 modulates the recruitment of SAR1s without GEF activity.

## Materials and Methods

### Plant material and growth conditions

The seeds of *Arabidopsis thaliana* wild-type (WT) (Columbia, Col-0) were obtained from the Arabidopsis Biological Resource Center (ABRC). Seeds of *A. thaliana* were surface-sterilized and germinated on half-strength modified Hoagland agar plates (Millner, 1992; Liu et al., 2012) and grown in the growth chamber at 22□ under a 16-h-light/8-h-dark cycle. The Pi-sufficient (+P) and Pi-deficient (−P) media were half-strength modified Hoagland nutrient solutions containing 250 μM and 0 μM KH_2_PO_4_, respectively. Seeds of *Nicotiana benthamiana* (*N. benthamiana*) were surface-sterilized and germinated on half-strength Murashige and Skoog medium and grown in the growth chamber at 24□ under a 16-h-light/8-h-dark cycle.

### Plasmid construction

For the self-complementing split-GFP assay (Liu et al., 2018), the constructs encoding *UBQ10:sXVE:*GFP1–10 (Addgene plasmid #125663), *UBQ10:sXVE:*ER-GFP1–10 (Addgene plasmid #125664) or *UBQ10:sXVE:*S11-GFP1–9 (Addgene plasmid #125665) was used. For the tripartite split-GFP association, the constructs encoding *UBQ10:sXVE:*GFP1–9 (Addgene plasmid #108187) or *35S:*GFP1–9.pMDC201 were used. The CDS of GFP1–9 was amplified by PCR using *P16*Δ*S:*GFP1–9 (Addgene plasmid #108186) as the template and subcloned into pMDC201, yielding *35S:*GFP1–9.pMDC201. For constructing the S10-tagged clones, the CDS of full-length and truncated *At*PHF1 and *At*PHT1;1 were amplified by PCR and subcloned into *UBQ10:XVE:*S10-(MCS) (Addgene plasmid #108177), in which the inducible promoter *UBQ10:sXVE* was replaced by the *UBQ10* promoter to generate *UBQ10:*S10-(MCS). The CDS of *At*SEC12 and *At*PHF1 were amplified by PCR and subcloned into *UBQ10:*S10-(MCS). For constructing the S11-tagged clones, the CDS of *At*PHT1;1 was amplified by PCR and subcloned into *UBQ10:sXVE:*(MCS)-3xHA-S11 (Addgene plasmid #108180). The CDS of *At*STP1 and *At*PHT1;1 and truncated *At*PHF1 were amplified by PCR and subcloned into *UBQ10:sXVE:*(MCS)-S11 (Addgene plasmid #108179), in which the inducible promoter *UBQ10:sXVE* was replaced by the *UBQ10* promoter to generate *UBQ10:*(MCS)-S11. The CDS of *At*SAR1b, *At*SAR1b[H74L] (Takeuchi, 2000), *At*SAR1b[T34N], *At*SAR1c, *At*SEC24a, *At*SEC13a, and *At*SEC16a were amplified and subcloned into *UBQ10:*(MCS)-S11. For constructing the GFP fusion clones, the inducible promoter *UBQ10:sXVE* of *UBQ10:sXVE:*(MCS)-GFP (Schlücking et al., 2013) was replaced by the *UBQ10* promoter to generate *UBQ10:*(MCS)-GFP. The CDS of *At*PHF1 was amplified and subcloned into *UBQ10:*(MCS)-GFP. For constructing the mCherry fusion clones, the GFP of *UBQ10:sXVE:*(MCS)-GFP (Schlücking et al., 2013) was replaced by mCherry to generate *UBQ10:sXVE:*(MCS)-mCherry. The CDS of *At*SEC12 was amplified and subcloned into *UBQ10:sXVE:*(MCS)-mCherry, followed by replacing the inducible promoter with the *UBQ10* promoter to generate *UBQ10:At*SEC12-mCherry. The CDS of mTb was amplified by PCR using R4pGWB601_UBQ10p-miniTurbo-NES-YFP (Addgene plasmid #127369) as the template and subcloned into *UBQ10:At*SAR1b-S11, yielding *UBQ10:At*SAR1b-mTb-S11. The abovementioned constructs were generated either by restriction enzyme cloning or by In-Fusion® HD cloning (Takara). For the yeast membrane-based split-ubiquitin assay, *At*PHT1;1-CubPLV, NubG-*At*CNIH5, and *At*OCT1-NubG constructs were obtained as previously described (Liu et al., 2024; Chiu et al., 2026). The CDS of *At*SAR1c were amplified by PCR, cloned into the pCR8/GW/TOPO, verified by sequencing. The *At*PHF1(noSTOP).pTOPO, cloned as previously described (Huang et al., 2013), and *At*SAR1c(noSTOP).pTOPO were recombined into XN22_GW (ABRC plasmid CD3-1735) via LR reaction to produce *At*PHF1-NubG and *At*SAR1c-NubG. Primer sequences and the restriction enzyme cutting sites used are listed in **Supplementary Table S1**.

### Agro-infiltration of *Nicotiana benthamiana* leaves

Three-to-four-week-old *Nicotiana benthamiana* plants were used for *A. tumefaciens* (strain EHA105)-mediated infiltration. The bacterial cultures were grown overnight in Luria-Bertani (LB) medium containing the appropriate antibiotics (50 μg/ml spectinomycin, 5 μg/ml rifampicin, or 50 μg/ml kanamycin) at 30°C with shaking at 200 rpm. Cells were pelleted and resuspended in infiltration buffer containing 10 mM MgCl_2_, 10 mM MES (pH 7.5), and 150 µM acetosyringone to an OD_600_ of 1.0 and incubated at room temperature (RT) for 2–4 h. Leaf infiltration was performed with agrobacterial suspensions at a final OD□□□ of 0.002–1.0. For *UBQ10:sXVE*-inducible expression, 9.18 μM β-estradiol was applied for an additional 24 or 48 h. Leaves were harvested at 48 or 72 h post-infiltration for confocal microscopy or protein extraction.

### Confocal image analysis

Confocal images were captured using the Zeiss LSM 800 microscope (Zeiss) equipped with Plan-Apochromat 10×/0.45 M27, 20×/0.8 M27, and 40×/1.3 Oil DIC M27 objectives. The acquisition was performed in multi-track mode with line switching, and the data were averaged over two readings. Excitation/emission wavelengths were 488 nm/410–546 nm and 561 nm/560–617 nm for GFP and mCherry, respectively. The z-stack was acquired with auto-optimized 50% overlapping optical sections spanning a total depth of 25–45 μm.

### Soluble and microsomal protein extraction

Soluble (S) and microsomal (M) fractions were prepared from agro-infiltrated *N. benthamiana* leaves with the modified low-speed pellet (LSP) method (Yoshimoto et al., 2004). Leaf tissues were ground in liquid nitrogen (N) and resuspended in three volumes (3 ml per gram tissue) of pre-chilled LSP buffer containing 100 mM Tris-HCl (pH 7.5), 400 mM sucrose, 1 mM EDTA, 1% Triton X-100, 0.1 mM phenylmethylsulfonyl fluoride (PMSF), and 1× Protease Inhibitor Cocktail (P9599, Sigma-Aldrich). After centrifugation at 500 × g for 5 min and then at 13,000 × g for 15 min at 4°C, the final supernatant was collected as the soluble fraction. The pellet was resuspended in pre-chilled LSP buffer lacking 1% Triton X-100 to yield the microsomal fraction.

### Proximity labeling assay

The *N. benthamiana* leaves were incubated in 50 µM biotin solution at RT for 30 min, ground with liquid N, and extracted in two volumes (2 ml per gram tissue) of lysis buffer containing 20 mM HEPES (pH 7.5), 40 mM KCl, 1 mM EDTA, 1% Triton X-100, 0.2 mM PMSF, and 1× Protease Inhibitor Cocktail (P9599, Sigma-Aldrich). After centrifugation at 4,000 × g for 5 min and then at 20,000 × g for 15 min at 4°C, the resulting supernatant was collected. A 25–750 µg of the crude extract was incubated with 5 µl streptavidin-agarose (S1638, Sigma-Aldrich) at 4□ overnight under 5 rpm end-to-end rotation. The beads collected by centrifugation at 2,500 × g and 4 □ for 5 min were washed with a lysis buffer containing 0.1% Triton X-100 once. After centrifugation at 2,500 × g for 5 min at 4□, the beads were eluted with 2× SDS sample buffer containing 100 mM Tris-HCl (pH 6.8), 20% glycerol, 4% SDS, and 100 mM dithiothreitol (DTT) with 0.4 M urea.

### Co-immunoprecipitation assay

The roots of 11-day-old Col-0, *UBQ10:*GFP, *UBQ10:At*SAR1c-GFP (Zeng et al., 2015), and *dex:At*SAR1b[H74L]-CFP (Lupanga et al., 2020) seedlings with or without 7 days of Pi starvation were collected. The seedlings of *dex:At*SAR1b[H74L]-CFP were treated with 10 µM dexamethasone (DEX) or DMSO for an additional 24 h before root collection. The roots were harvested for total protein extraction with lysis buffer containing 25 mM HEPES (pH 7.2), 150 mM NaCl, 0.5% IGEPAL CA-630 (I8896, Sigma-Aldrich), 2 mM EDTA, 0.5 mM MgCl_2,_ and 1× Protease Inhibitor Cocktail (P9599, Sigma-Aldrich). After centrifugation at 16,000 × g for 5 min at 4°C, supernatants were cross-linked with 2.5 mM dithiobis [succinimidyl propionate] (DSP) at RT for 30 min and quenched with 100 mM Tris (pH 7.4) for 15 min. A 15–750 µg of protein mixture was incubated with 5 µl of GFP-Trap® Agarose (ChromoTek) at 4□ for 2 h with end-over-end rotation at 5 rpm, washed with the lysis buffer once, centrifuged at 2,500 × g at 4□ for 5 min, and eluted with 2× SDS sample buffer containing 100 mM Tris-HCl (pH 6.8), 20% glycerol, 4% SDS, and 100 mM DTT.

### Immunoblot analysis

Protein samples were loaded onto 4–12% Q-PAGE™ Bis-Tris Precast Gel (SMOBIO) and transferred to polyvinylidene difluoride (PVDF) membranes (IPVH00010 Immobilon-P Membrane, Merck). The membrane was blocked with 1% BSA in 1× PBS solution containing 0.2% Tween 20 (PBST, pH 7.2) at RT for 1 h and hybridized with primary antibodies: anti-GFP (1:5,000, AS21 4696; Agrisera) at 4□ for overnight, anti-*At*PHT1;1/2/3 (1:1,000, PHY0753; PhytoAB) at 4□ for overnight, anti-*At*PHF1 (Huang et al., 2013) at RT for 2 h, anti-PIP2;7 (1:5,000, AS22 4812, Agrisera) at RT for 1 h, anti-cFBPase (1:5000, AS04 043, Agrisera) at 4□ for overnight, anti-mCherry (1:1,000, ab167453, Abcam) at RT for 1 h, and anti-S11 (1:10,000) at RT for 1 h, the anti-S11 polyclonal rabbit antibody was raised against the peptide of S11 (EKRDHMVLLEYVTAAGITDASC) and produced by LTK BioLaboratories, Taiwan. The membrane was washed four times with 1× PBST for 5 min, followed by hybridization with the horseradish peroxidase-conjugated secondary antibody (1:20,000–1:40,000, GTX213110-01 and 1:10,000, GTX213111-01, GeneTex) in blocking solution at RT for 1 h. After four washes in 1× PBST for 5 min and a rinse with distilled water, chemiluminescent substrates (WesternBright^TM^ ECL and WesternBright^TM^ Sirius, Advansta) for signal detection were applied.

### Quantification analyses

For puncta quantification, two non-overlapping regions of interest (ROIs, 1000 µm²) were randomly selected from a single confocal image (53.24 × 53.24 µm²). For each ROI, the automatic Triangle threshold was applied. Punctate structures were defined using the ImageJ (Schneider et al., 2012) ‘analyze particles’ function with circularity from 0.1 to 1.0 and size from 0.04 to infinity. Co-localization analysis was performed with a Costes’ automatic threshold, and Manders’ overlap coefficients were calculated using the JACoP plugin with object-based method in Fiji (Bolte and Cordelieres, 2006). The Dunnett’s test and Mann-Whitney U test were performed using JASP 0.95.4.0. Puncta diameters were measured by using the Zeiss ZEN 3.7 software. The box plots were generated by using BoxPlotR (Spitzer et al., 2014).

### Chemical treatment

Acetosyringone (150 mM stock in DMSO, D134406, Sigma-Aldrich) was diluted to a final concentration of 150 µM in ddH_2_O. β-estradiol (36.7 mM stock in ethanol, E2758, Sigma-Aldrich) was diluted to a final concentration of 9.18 μM in ddH_2_O. Dithiobis [succinimidyl propionate] (25 mM stock in DMSO, 22586, ThermoFisher) was diluted to a final concentration of 2.5 mM in ddH_2_O. Dexamethasone (100 mM stock in DMSO, D4902, Sigma-Aldrich) was diluted to a final concentration of 10 µM in ddH_2_O.

### Accession Numbers

Sequence data from this article can be found in the Arabidopsis Genome Initiative under the following accession numbers: *At*PHF1 (AT3G52190), *At*SEC12 (AT2G01470), *At*PHT1;1 (AT5G43350), *At*STP1 (AT1G11260), *At*SAR1b (AT1G56330), *At*SAR1c (AT4G02080), *At*SEC24a (AT3G07100), *At*SEC13a (AT3G01340), *At*SEC16a (AT5G47480), *At*CNIH5 (AT4G12090), *At*OCT1 (AT1G73220).

## Supplementary Data

**Supplementary Fig. S1. Expression of ER lumen marker in the absence and presence of *At*PHT1;1 and *At*PHF1 overexpression.** Representative z-stack confocal images **(A)** and quantification of puncta numbers **(B)** of the ER-luminal marker mCherry-HDEL expressed alone or co-expressed with *At*PHF1-GFP and/or *At*PHT1;1-S11. Box plots show medians (center lines), interquartile ranges (boxes), and data ranges (whiskers), along with individual data points (dots). The number of regions of interest (ROI) used for quantification is shown in parentheses; the data were collected from three independent experiments. Not significant, n.s.; Dunnett’s test for multiple comparisons against mCherry-HDEL expressed alone. Scale bars, 5 µm.

**Supplementary Fig. S2. Size analysis of *At*PHF1-GFP- and mCherry-HDEL-labeled puncta in the presence of *At*PHT1;1-S11 overexpression. (A)** Quantification of puncta number upon overexpression with *At*PHT1;1-S11, corresponding to Fig. 1D. Data, shown as mean ± standard error (SE), were collected from three independent experiments (Exp. 1, Exp. 2, and Exp. 3). n = the number of region of interest (ROI) counted. **(B)** Size distribution of overlapping and non-overlapping puncta labeled by *At*PHF1-GFP or the ER luminal marker mCherry-HDEL, corresponding to (**A**). Box plots show medians (center lines), interquartile ranges (boxes), and data ranges (whiskers), along with individual data points (dots). The number of puncta used for quantification is shown in parentheses. Dunnett’s test for multiple comparisons against overlapping puncta. Not significant, n.s..

**Supplementary Fig. S3. Expression of *AtSAR1* isoforms in Col-0 under Pi deprivation by RNA-seq analysis.** Expression of *AtSAR1a/b/c/d/e* in the shoot and root of 10-day-old WT (Col-0) seedlings under Pi-sufficient conditions (+P0) or under one day (–P1) and three days of Pi starvation (–P3) as previously described (Liu et al., 2016). RPKM stands for reads per kilobase of transcript per million mapped reads. Numbers represent average RPKM values of two replicates, with an assigned value of 0.25 for readings below this threshold.

**Supplementary Fig. S4. The expression and distribution of COPII-related components in agro-infiltrated *N. benthamiana* leaves. (A)** Expression and distribution of S11-tagged *At*SAR1b, *At*SAR1c, *At*SEC24a, *At*SEC13a, and *At*SEC16a at the peripheral layer and 3xHA-S11 at the middle layer of agro-infiltrated tobacco epidermal cells. **(B)** Expression and distribution of S10-tagged *At*SEC12, *At*PHF1, and 3xHA at the middle layer of agro-infiltrated tobacco epidermal cells. Scale bars, 5 µm. Representative confocal images are shown from three independent experiments.

**Supplementary Fig. S5. Quantification of puncta sizes upon co-expression of *At*PHF1-mCherry and *At*PHT1;1 with ERES markers. (A, B)** Quantification of puncta number upon overexpression with *At*SAR1b-S11 (**A**) or *At*SEC24a-S11 (**B**), corresponding to Fig. 2C. Data, shown as mean ± standard error (SE), were collected from three independent experiments (Exp. 1, Exp. 2, and Exp. 3). n = the number of region of interest (ROI) counted. **(C**, **D)** Size distribution of overlapping and non-overlapping puncta labeled by *At*PHF1-mCherry or ERES markers, corresponding to (**A**) and (**B**), respectively. Box plots show medians (center lines), interquartile ranges (boxes), and data ranges (whiskers), along with individual data points (dots). The number of puncta used for quantification is shown in parentheses. Dunnett’s test for multiple comparisons against overlapping puncta. Not significant, n.s..

**Supplementary Fig. S6. Imaging analysis of ERES marker in the absence and presence of *At*PHT1;1 and/or *At*PHF1 overexpression. (A)** Expression and distribution of *At*SAR1b-S11 in the absence and presence of *At*PHF1-mCherry and *At*PHT1;1. Representative confocal images at the peripheral layers of the epidermis are shown. Scale bars, 5 µm. **(B)** Quantification analysis of *At*SAR1b-labeled ERES puncta of (**A**) from three independent experiments. The number of regions of interest (ROI) used for quantification is shown in parentheses. Box plots show medians (center lines), interquartile ranges (boxes), and data ranges (whiskers), along with individual data points (dots). Not significant, n.s.; ***, *P* < 0.001; Dunnett’s test for multiple comparisons against ERES marker expressed alone.

**Supplementary Fig. S7. The interaction analysis of *At*PHF1 and *At*SAR1b by proximity labeling.** Thirty times more leaf total protein extract from *N. benthamiana* expressing *At*SAR1b-mTb-S11 was utilized for streptavidin pulldown compared to the extract expressing mTb-NES-EYFP. Anti-S11 antibody was used to detect EYFP-tagged and S11-tagged fusion proteins. Representative results are shown from two independent experiments.

**Supplementary Fig. S8. Expression and subcellular distribution of the truncated *At*PHF1 variants in agro-infiltrated *N. benthamiana* leaves. (A)** Schematic diagrams of the truncated *At*PHF1 variants with domain boundaries indicated by amino acid numbers. **(B, C)** The expression and distribution of the S10-tagged (**B**) and the S11-tagged (**C**) truncated *At*PHF1 variants at the middle layer of epidermal cells. Scale bars, 20 µm. Representative confocal images are shown from three independent experiments.

**Supplementary Fig. S9. *At*SAR1b and *At*SAR1c interact with the cytosolic and transmembrane domains of *At*PHF1 in agro-infiltrated *N. benthamiana* leaves.** (A–C) The interaction analyses of the S10-*At*PHF1 truncated variants with *At*SAR1b-S11 (**A**), *At*SAR1c-S11 (**B**), and 3xHA-S11 (**C**) by using the tripartite split-GFP assay. The interactions with 3xHA-S11 were used as negative controls. Scale bars, 10 µm. Representative confocal images taken at the peripheral layer of the epidermis are shown from at least three independent experiments.

**Supplementary Fig. S10. Interaction analysis of *At*SAR1c-GFP with endogenous *At*PHF1 and *At*PHT1;1/2/3.** Eleven-day-old wild-type (Col-0), *UBQ10:*GFP, and *UBQ10:At*SAR1c-GFP seedlings were subjected to Pi deprivation (0 µM KH_2_PO_4_, 7 days of starvation). Fifty-fold more root protein extract expressing *At*SAR1c-GFP was used as input for GFP-Trap co-immunoprecipitation relative to that expressing GFP. Anti-S11 antibody was used to detect GFP fusion proteins. Representative results are shown from three independent experiments.

**Supplementary Fig. S11. Interaction analysis of *At*PHF1 with *At*SAR1 in the yeast split-ubiquitin system. (A)** Co-expression of *At*PHF1-CubPLV with NubG-*At*PHT1;1, *At*SAR1b-NubG or *At*SAR1c-NubG. **(B)** Co-expression of *At*SEC12-CubPLV with *At*SAR1b-NubG and *At*SAR1c-NubG. The co-expression of *At*PHF1-CubPLV or *At*SEC12-CubPLV with NubWT was used as a positive control, while the co-expression with NubG was used as a negative control. The CDS of *At*PHF1, *At*SEC12, *At*PHT1;1, *At*SAR1b, and *At*SAR1c were amplified by PCR, cloned into the pCR8/GW/TOPO, verified by sequencing, and recombined into destination vectors via LR reactions. *At*PHF1 and *At*SEC12 were cloned into synthesized MET17-PLVCub_GW to generate C-terminal Protein A–LexA–VP16–Cub fusions (*At*PHF1-CubPLV and *At*SEC12-CubPLV); *At*PHT1;1 was cloned into NX32_GW (ABRC plasmid CD3-1737) to produce NubG-*At*PHT1;1; *At*SAR1b and *At*SAR1c were cloned into XN22_GW (ABRC plasmid CD3-1735) to produce *At*SAR1b-NubG and *At*SAR1c-NubG. Yeast transformants were grown on synthetic medium lacking tryptophan and leucine (SC–WL) for growth detection or on synthetic medium lacking tryptophan, leucine, adenine, and histidine containing 0.5 µM methionine (SC–WLAH + M) for testing protein-protein interaction. Representative results are shown from two independent experiments.

**Supplementary Fig. S12. Distribution of *At*SAR1b/[H74L]/[T34N] in the agro-infiltrated *N. benthamiana* leaves. (A)** The expression and distribution of the S11-tagged GTP-locked ([H74L]) and GDP-locked *At*SAR1b ([T34N]) at the peripheral layer of epidermal cells. Scale bars, 5 µm. Representative confocal images are shown from three independent experiments. **(B)** Protein expression of *At*SAR1b/[H74L]/[T34N] in the soluble (S) and microsomal (M) fractions isolated using the modified low-speed pellet (LSP) method (Yoshimoto et al., 2004). Anti-S11 antibody was used to detect S11 fusion proteins. The membrane association percentage (%) was calculated as M/(S+M) and is shown as mean ± standard error (SE) from two independent experiments. Aquaporin PIP2;7 and cytosolic fructose-1,6-bisphosphatase (cFBPase) served as microsomal and soluble controls, respectively. Amido black staining was used as a loading control.

**Supplementary Fig. S13. ER export-incompetent *At*PHT1;1 overexpression cannot trigger partial distribution of *At*PHF1 to punctate structures in agro-infiltrated *N*. *benthamiana* leaves. (A)** Expression and distribution of S10-*At*PHT1;1 TM1–6 and *At*PHT1;1 TM7–12-S11 upon co-expression with *At*PHF1-mCherry in agro-infiltrated tobacco epidermal cells. Scale bars, 5 µm. Representative confocal images taken at the peripheral layer of epidermal cells are shown from three independent experiments. **(B)** Distribution of *At*PHF1-GFP in the presence of S10-*At*PHT1;1 TM1–6 or *At*PHT1;1 TM7–12-S11 co-expression. Scale bars, 5 µm. Representative confocal images are shown from three independent experiments. **(C)** Quantification of *At*PHF1-GFP-labeled punctate structures in (**B**). The data for *At*PHF1-GFP expressed alone are the same as those shown in Fig. 1B. Box plots display medians (center lines), interquartile ranges (boxes), and data ranges (whiskers), along with individual data points (dots). The number of regions of interest used for quantification is shown in parentheses. Dunnett’s test for multiple comparisons against *At*PHF1-GFP expressed alone. Not significant, n.s..

**Supplementary Fig. S14. Interaction analysis of the N-terminal cytosolic domain of *At*PHF1 with *At*SAR1b/[H74L]/[T34N] in the yeast two-hybrid assay.** GAL4[BD]-S10-*At*PHF1[N was co-expressed with GAL4[AD]-*At*SAR1b-S11, GAL4[AD]-*At*SAR1b[H74L]-S11, or GAL4[AD]-*At*SAR1b[T34N]-S11. The co-expression of Krev1 with RalGDS-wt, RalGDS-m1, and RalGDS-m2 were strong positive, weak positive, and negative controls, respectively. The CDS of *At*SAR1b-S11, *At*SAR1b[H74L]-S11, and *At*SAR1b[T34N]-S11 were subcloned into pCR8/GW/TOPO and recombined into pDEST22 via LR reaction to generate GAL4[AD]-*At*SAR1b/[H74L]/[T34N]-S11; the CDS of S10-*At*PHF1 N was subcloned into pCR8/GW/TOPO and recombined into pDEST32 via LR reaction to generate GAL4[BD]-S10-*At*PHF1 N. Yeast transformants were grown on synthetic medium lacking tryptophan and leucine (SC–WL) for growth detection, and on medium lacking tryptophan, leucine, and uracil (SC–WLU) or lacking tryptophan, leucine, and histidine with 10 mM 3-Amino-1,2,4-triazole (SC–WLH + 3AT) to assess protein-protein interactions. Representative results are shown from two independent experiments.

**Supplementary Table S1. Oligonucleotides used for plasmid constructs.**

## Acknowledgments

We thank Dr. Liwen Jiang at the Chinese University of Hong Kong, People’s Republic of China, for sharing the *Arabidopsis* seeds of *UBQ10:At*SAR1c-GFP and *UBQ10:*GFP homozygous lines, Dr. Masaki Takeuchi at the University of Tokyo, Japan, for sharing the clone *35S:At*SAR1b[H74L], Dr. Tzyy-Jen Chiou at Academia Sinica, Taiwan (R.O.C.), for sharing the clone *35S:At*PHF1-mCherry, and Dr. Karin Schumacher at the Cell Biology Research Group of the Centre for Organismal Studies (COS), Heidelberg University, Germany, for sharing the *Arabidopsis* seeds of *dex:At*SAR1b[H74L]-CFP T2 lines. We acknowledge Ms. Wen-Chun Chou for constructing the plasmid encoding *35S:*GFP1–10 and the technical support from the Confocal Image Core, National Tsing Hua University (sponsored by MOST 108-2731-M-007-001 and MOST 110-2731-M-007-001).

## Author Contributions

T.-Y. L. conceived the original research plan, designed and supervised the experiments, and wrote the article. H.-F. L. designed and performed the experiments, analyzed the data, wrote the article, and contributed to the preparation of figures. J.-D. C. designed and performed the experiments.

## Data Availability Statement

All study data are included in the article and/or Supporting Material.

