## Supplemental Figures and Table for "Involvement of PHOSPHATE TRANSPORTER TRAFFIC FACILITATOR1 in COPII assembly by interacting with SAR1 GTPase"

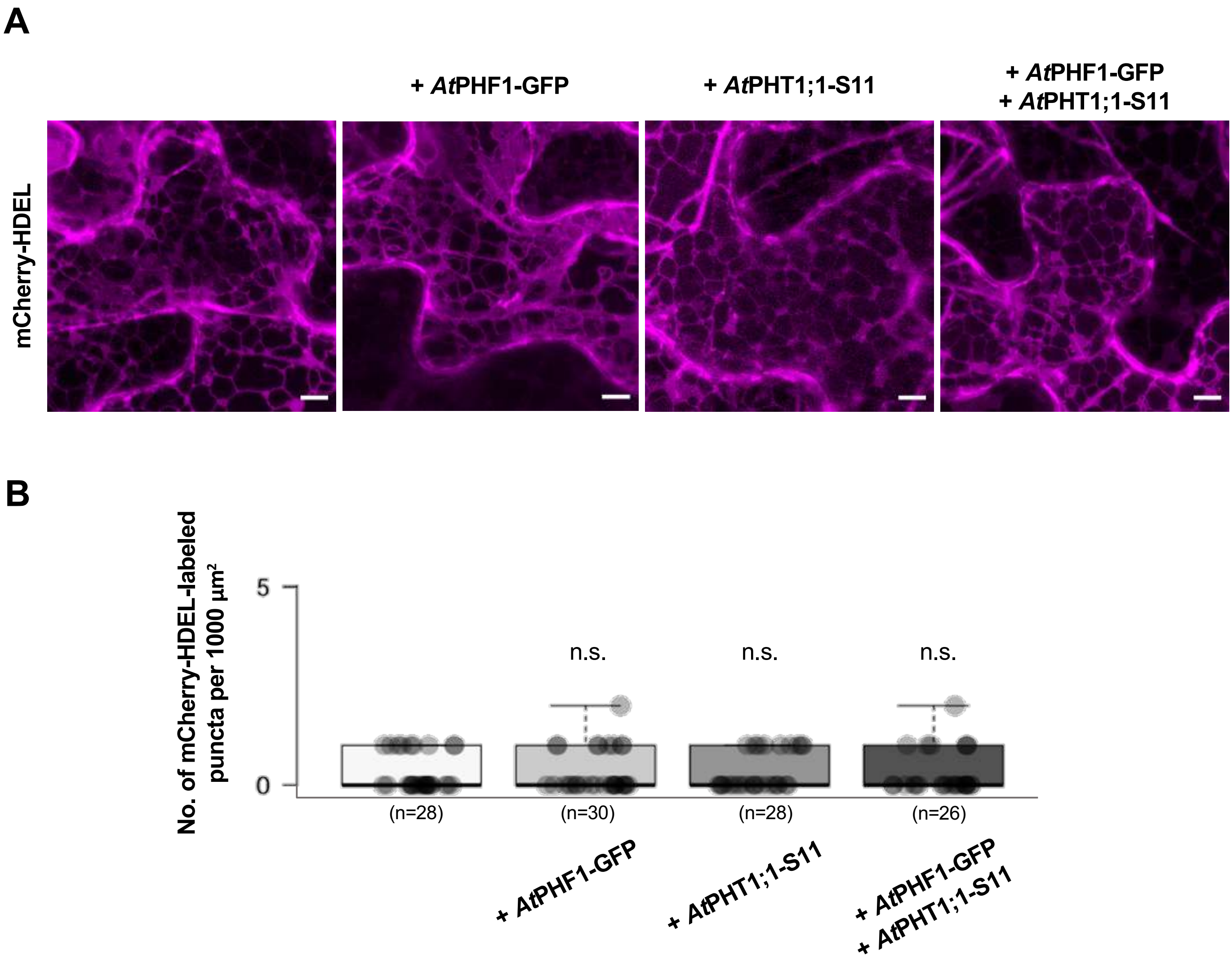


**Supplementary Fig. S1. Expression of ER lumen marker in the absence and presence of *At*PHT1;1 and *At*PHF1 overexpression.** Representative z-stack confocal images (**A**) and quantification of puncta numbers (**B**) of the ER-luminal marker mCherry-HDEL expressed alone or co-expressed with *At*PHF1-GFP and/or *At*PHT1;1-S11. Box plots show medians (center lines), interquartile ranges (boxes), and data ranges (whiskers), along with individual data points (dots). The number of regions of interest (ROI) used for quantification is shown in parentheses; the data were collected from three independent experiments. Not significant, n.s.; Dunnett's test for multiple comparisons against mCherry-HDEL expressed alone. Scale bars, 5 µm.

**A**

| | No. of ROI with puncta / No. of ROI counted | Number of puncta per 1000 μm² | | |
|---|---|---|---|---|
| | | Overlapping puncta | Non-overlapping puncta | |
| | | | mCherry-HDEL alone | *At*PHF1-GFP alone |
| Exp. 1 | 16/18 | 0.5±0.3 | 0.9±0.6 | 3.3±1.6 |
| Exp. 2 | 6/10 | 1.5±1.0 | 2.0±1.2 | 12.3±1.8 |
| Exp. 3 | 8/12 | 0.2±0.2 | 1.7±0.9 | 2.3±0.9 |

**B**

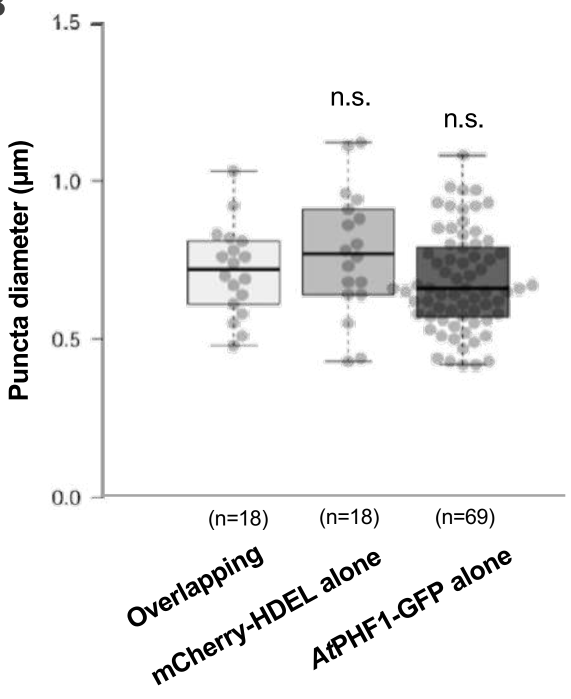


**Supplementary Fig. S2. Size analysis of *At*PHF1-GFP- and mCherry-HDEL-labeled puncta in the presence of *At*PHT1;1-S11 overexpression.** (**A**) Quantification of puncta number upon overexpression with *At*PHT1;1-S11, corresponding to Fig. 1D. Data, shown as mean ± standard error (SE), were collected from three independent experiments (Exp. 1, Exp. 2, and Exp. 3). n = the number of region of interest (ROI) counted. (**B**) Size distribution of overlapping and non-overlapping puncta labeled by *At*PHF1-GFP or the ER luminal marker mCherry-HDEL, corresponding to (**A**). Box plots show medians (center lines), interquartile ranges (boxes), and data ranges (whiskers), along with individual data points (dots). The number of puncta used for quantification is shown in parentheses. Dunnett's test for multiple comparisons against overlapping puncta. Not significant, n.s..

| | | Shoot | | | Root | | |
|---|---|---|---|---|---|---|---|
| **AGI** | **Gene Name** | **+P0** | **–P1** | **–P3** | **+P0** | **–P1** | **–P3** |
| AT1G09180 | *At*SAR1a | 0.47 | 0.85 | 1.17 | 0.43 | 0.51 | 1.13 |
| AT1G56330 | *At*SAR1b | 87.20 | 81.73 | 74.61 | 157.96 | 181.61 | 190.43 |
| AT4G02080 | *At*SAR1c | 85.28 | 71.46 | 76.11 | 128.42 | 133.27 | 132.56 |
| AT3G62560 | *At*SAR1d | 35.26 | 33.74 | 35.03 | 47.56 | 48.86 | 47.94 |
| AT1G02620 | *At*SAR1e | 9.64 | 2.21 | 2.35 | 0.25 | 0.25 | 0.25 |

**Supplementary Fig. S3. Expression of *AtSAR1* isoforms in Col-0 under Pi deprivation by RNA-seq analysis.** Expression of *AtSAR1a/b/c/d/e* in the shoot and root of 10-day-old WT (Col-0) seedlings under Pi-sufficient conditions (+P0) or under one day (–P1) and three days of Pi starvation (–P3) as previously described (Liu et al., 2016). RPKM stands for reads per kilobase of transcript per million mapped reads. Numbers represent average RPKM values of two replicates, with an assigned value of 0.25 for readings below this threshold.

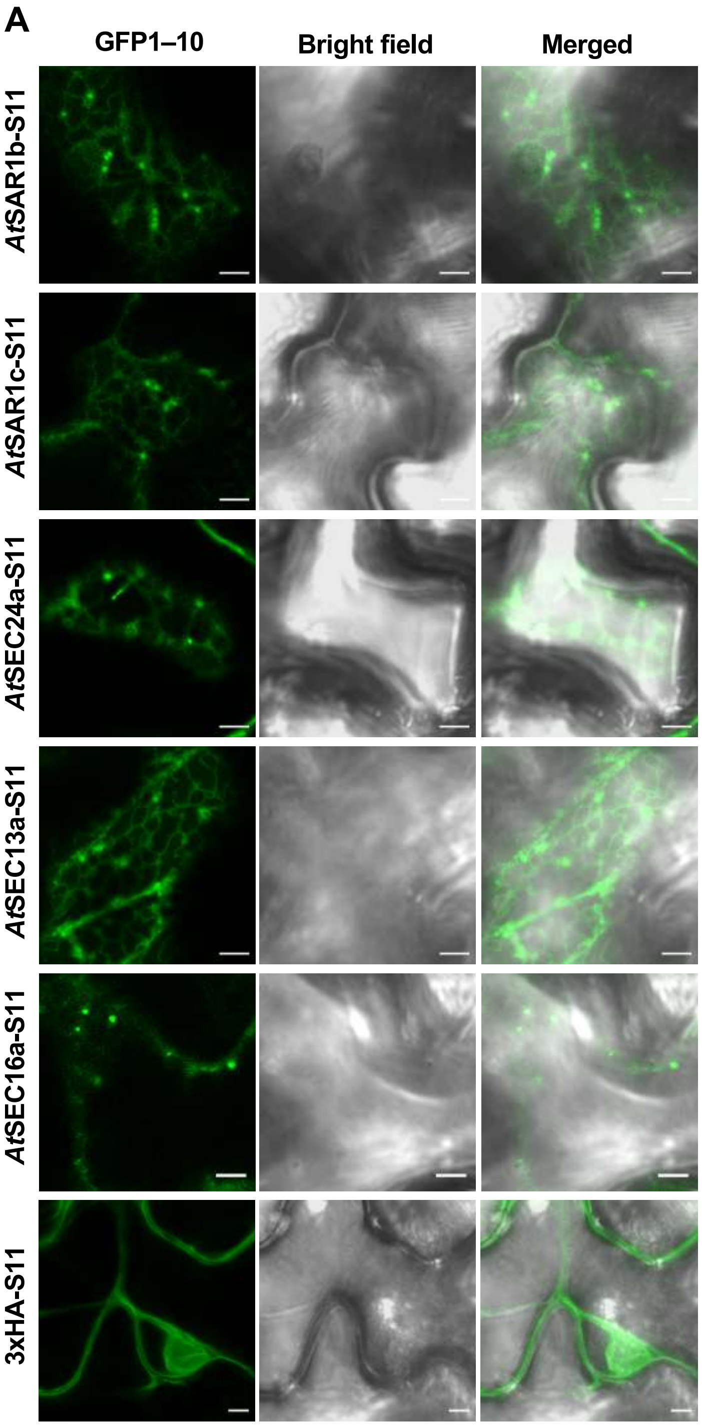

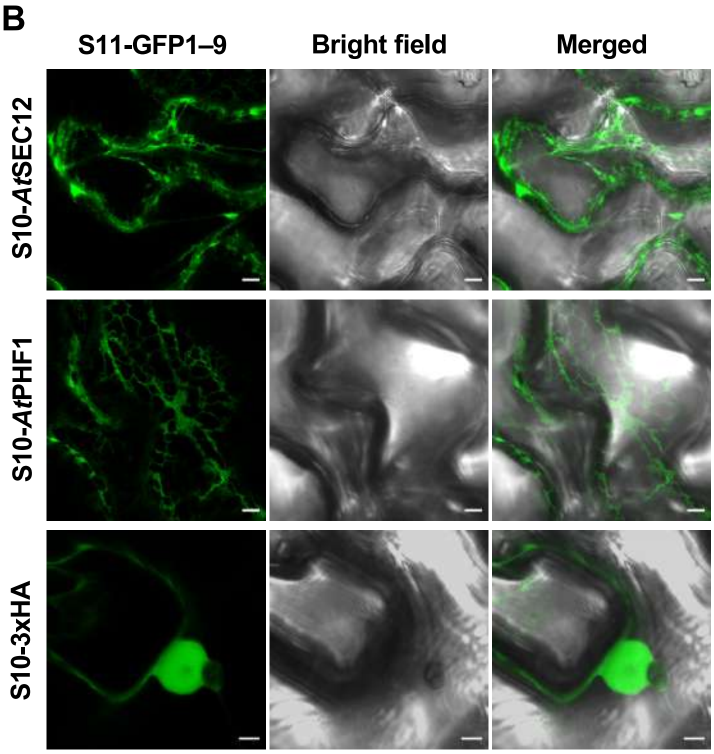


**Supplementary Fig. S4. The expression and distribution of COPII-related components in agro-infiltrated *N. benthamiana* leaves.** (**A**) Expression and distribution of S11-tagged *At*SAR1b, *At*SAR1c, *At*SEC24a, *At*SEC13a, and *At*SEC16a at the peripheral layer and 3xHA-S11 at the middle layer of agro-infiltrated tobacco epidermal cells. (**B**) Expression and distribution of S10-tagged *At*SEC12, *At*PHF1, and 3xHA at the middle layer of agro-infiltrated tobacco epidermal cells. Scale bars, 5 µm. Representative confocal images are shown from three independent experiments.

**A**

| | No. of ROI with puncta / No. of ROI counted | Number of puncta per 1000 μm² | | |
|---|---|---|---|---|
| | | Overlapping puncta | Non-overlapping puncta | |
| | | | *At*SAR1b alone | *At*PHF1-mCherry alone |
| Exp. 1 | 6/10 | 0.5±0.3 | 1.2±0.5 | 0.3±0.2 |
| Exp. 2 | 7/10 | 0.8±0.3 | 0.4±0.3 | 0.9±0.1 |
| Exp. 3 | 8/8 | 0.8±0.4 | 0.4±0.2 | 1.1±0.3 |

**B**

| | No. of ROI with puncta / No. of ROI counted | Number of puncta per 1000 μm² | | |
|---|---|---|---|---|
| | | Overlapping puncta | Non-overlapping puncta | |
| | | | *At*SEC24a alone | *At*PHF1-mCherry alone |
| Exp. 1 | 5/10 | 1.0±0.4 | 0.0±0.0 | 0.4±0.2 |
| Exp. 2 | 10/12 | 1.1±0.4 | 0.7±0.3 | 0.8±0.3 |
| Exp. 3 | 10/10 | 1.3±0.4 | 0.7±0.3 | 0.4±0.2 |

**C**

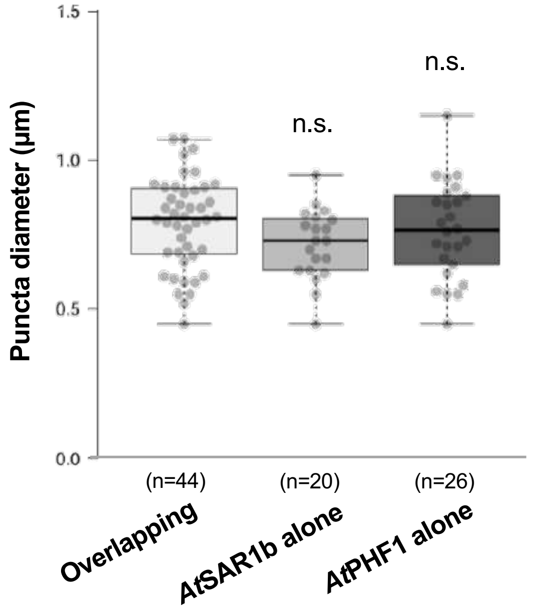


**D**

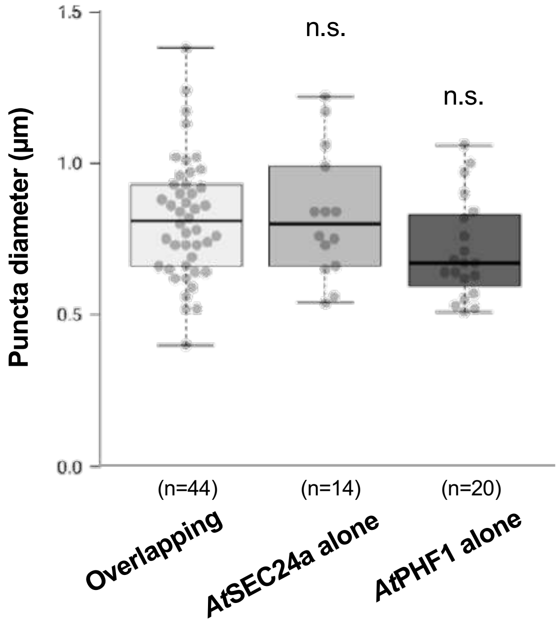


**Supplementary Fig. S5. Quantification of puncta sizes upon co-expression of *At*PHF1-mCherry and *At*PHT1;1 with ERES markers. (A, B)** Quantification of puncta number upon overexpression with *At*SAR1b-S11 (**A**) or *At*SEC24a-S11 (**B**), corresponding to Fig. 2C. Data, shown as mean ± standard error (SE), were collected from three independent experiments (Exp. 1, Exp. 2, and Exp. 3). n = the number of region of interest (ROI) counted. **(C, D)** Size distribution of overlapping and non-overlapping puncta labeled by *At*PHF1-mCherry or ERES markers, corresponding to (**A**) and (**B**), respectively. Box plots show medians (center lines), interquartile ranges (boxes), and data ranges (whiskers), along with individual data points (dots). The number of puncta used for quantification is shown in parentheses. Dunnett's test for multiple comparisons against overlapping puncta. Not significant, n.s..

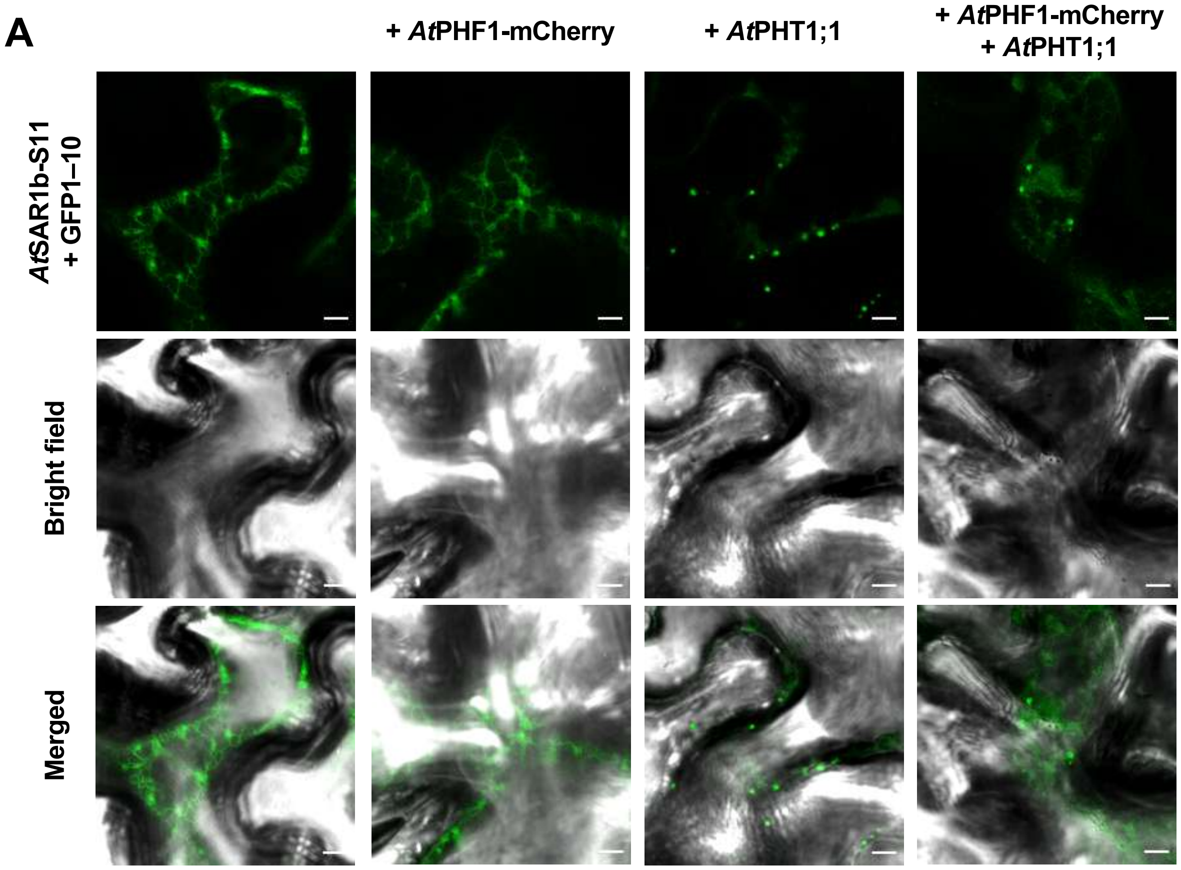


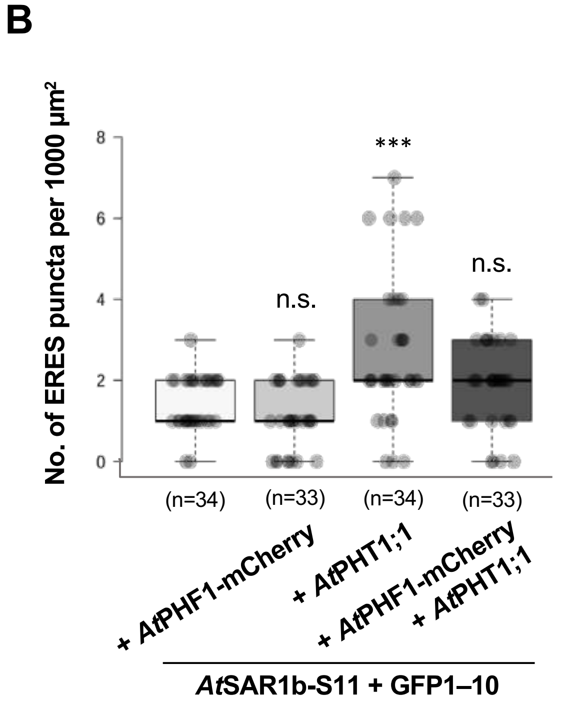


**Supplementary Fig. S6. Imaging analysis of ERES marker in the absence and presence of *At*PHT1;1 and/or *At*PHF1 overexpression.** (**A**) Expression and distribution of *At*SAR1b-S11 in the absence and presence of *At*PHF1-mCherry and *At*PHT1;1. Representative confocal images at the peripheral layers of the epidermis are shown. Scale bars, 5 µm. (**B**) Quantification analysis of *At*SAR1b-labeled ERES puncta of (**A**) from three independent experiments. The number of regions of interest (ROI) used for quantification is shown in parentheses. Box plots show medians (center lines), interquartile ranges (boxes), and data ranges (whiskers), along with individual data points (dots). Not significant, n.s.; ***, $P < 0.001$; Dunnett's test for multiple comparisons against ERES marker expressed alone.

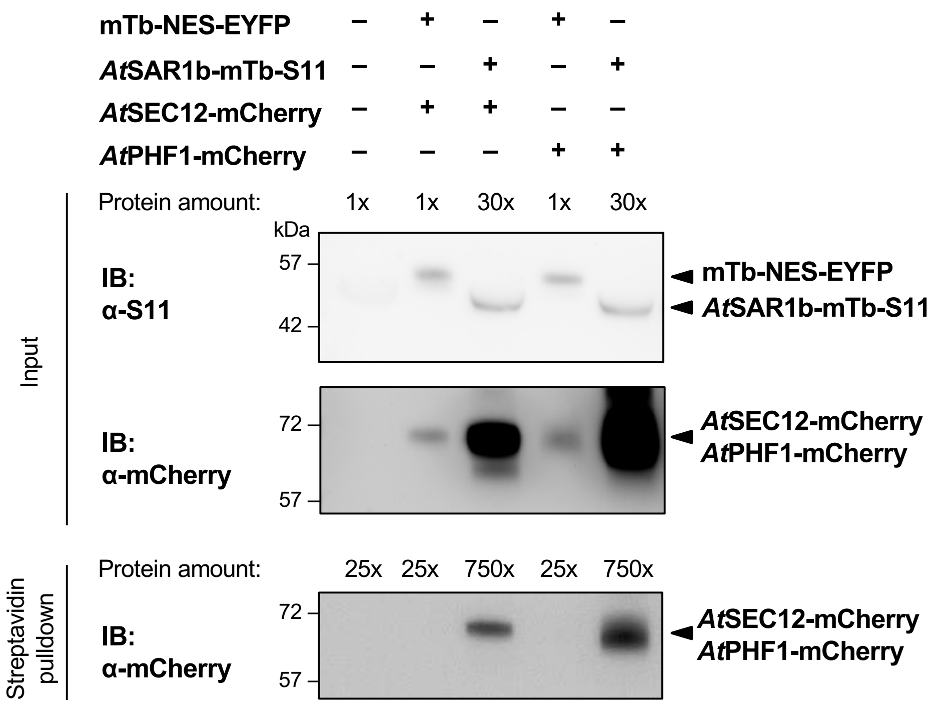


**Supplementary Fig. S7. The interaction analysis of *At*PHF1 and *At*SAR1b by proximity labeling.** Thirty times more leaf total protein extract from *N. benthamiana* expressing *At*SAR1b-mTb-S11 was utilized for streptavidin pulldown compared to the extract expressing mTb-NES-EYFP. Anti-S11 antibody was used to detect EYFP-tagged and S11-tagged fusion proteins. Representative results are shown from two independent experiments.

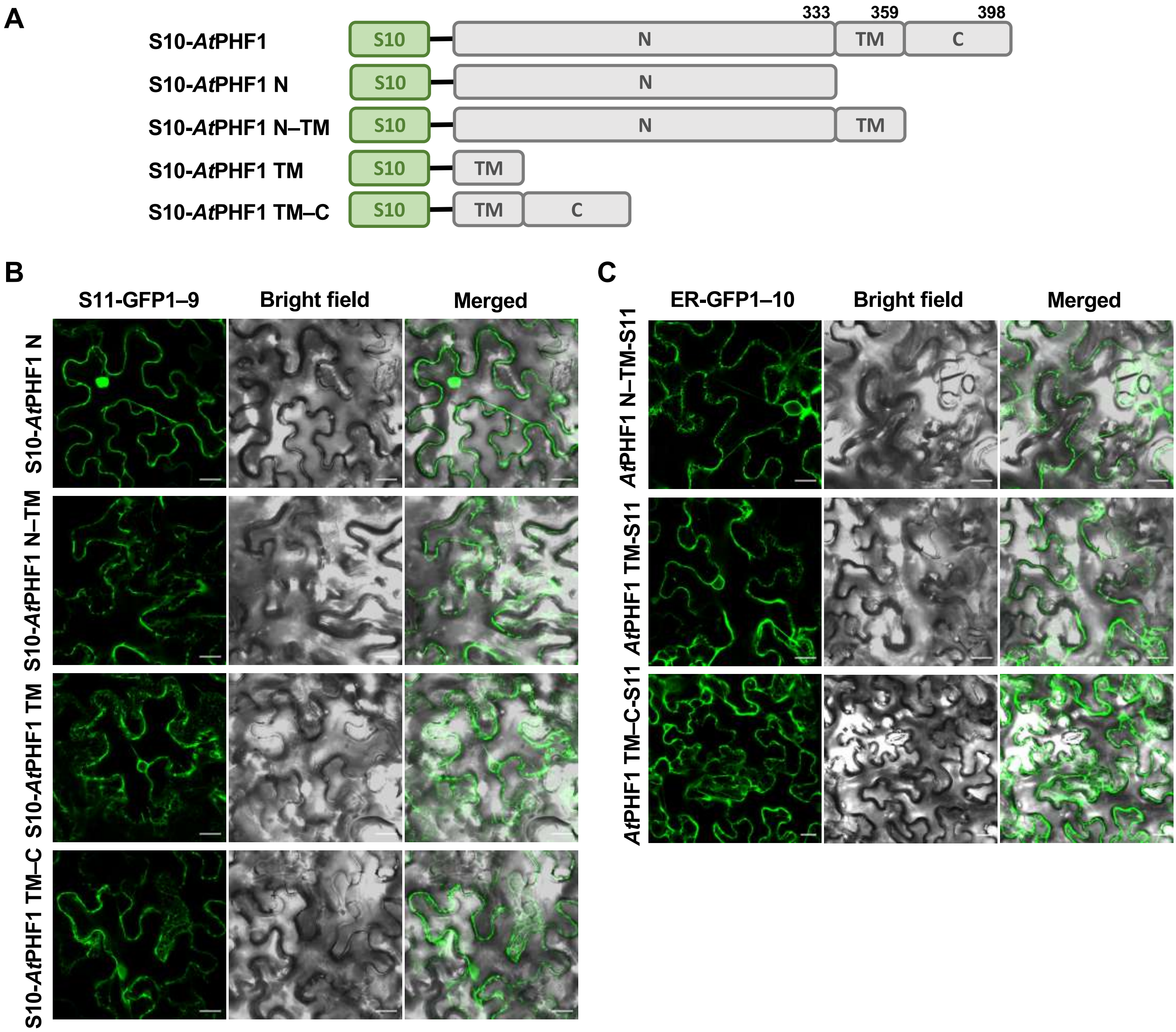


**Supplementary Fig. S8. Expression and subcellular distribution of the truncated *At*PHF1 variants in agro-infiltrated *N. benthamiana* leaves.** (**A**) Schematic diagrams of the truncated *At*PHF1 variants with domain boundaries indicated by amino acid numbers. (**B**, **C**) The expression and distribution of the S10-tagged (**B**) and the S11-tagged (**C**) truncated *At*PHF1 variants at the middle layer of epidermal cells. Scale bars, 20 µm. Representative confocal images are shown from three independent experiments.

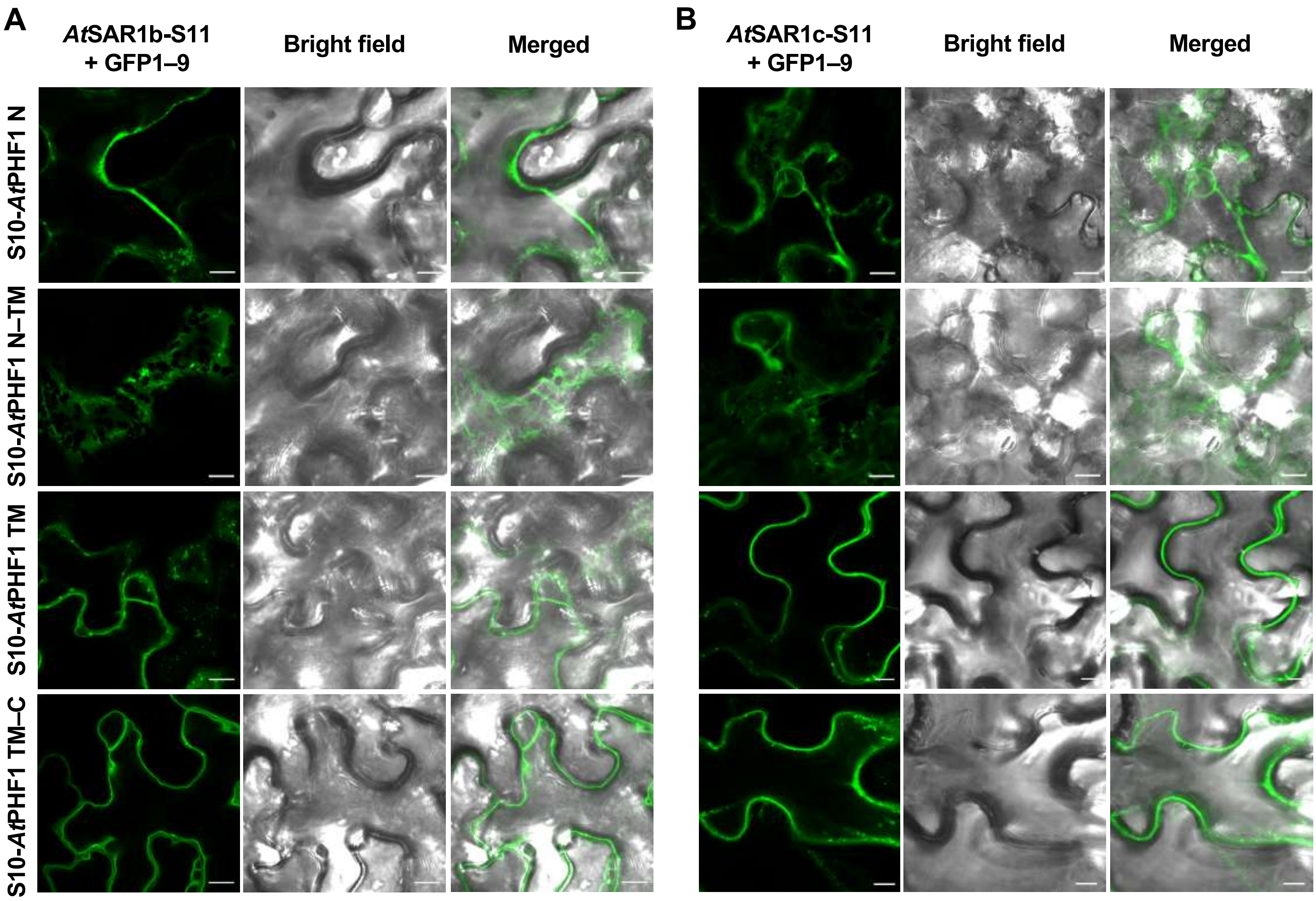


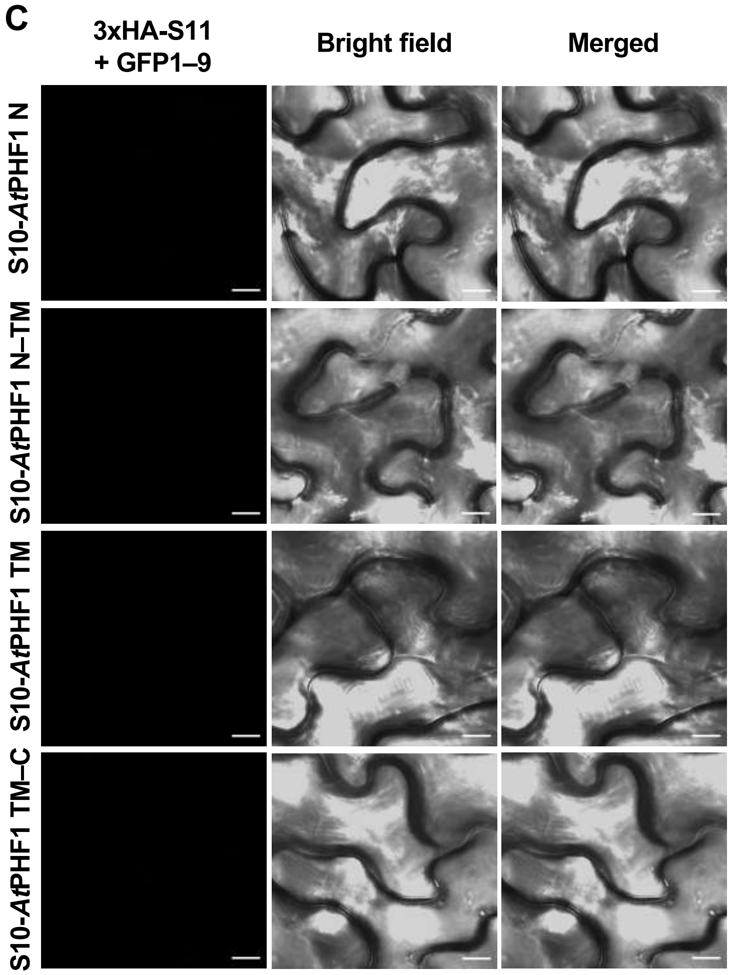


**Supplementary Fig. S9. *At*SAR1b and *At*SAR1c interact with the cytosolic and transmembrane domains of *At*PHF1 in agro-infiltrated *N. benthamiana* leaves.** (**A**–**C**) The interaction analyses of the S10-*At*PHF1 truncated variants with *At*SAR1b-S11 (**A**), *At*SAR1c-S11 (**B**), and 3xHA-S11 (**C**) by using the tripartite split-GFP assay. The interactions with 3xHA-S11 were used as negative controls. Scale bars, 10 μm. Representative confocal images taken at the peripheral layer of the epidermis are shown from at least three independent experiments.

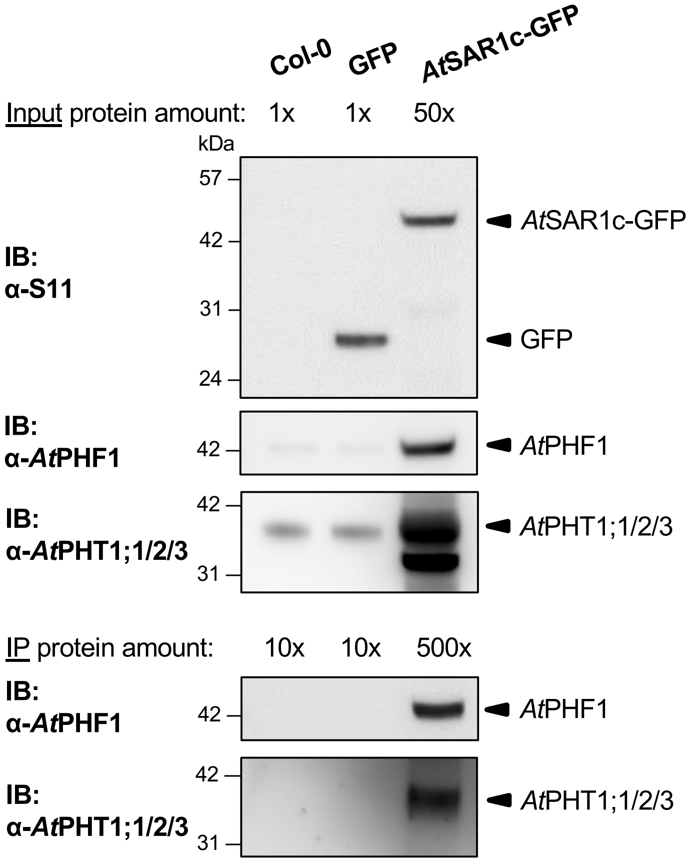


**Supplementary Fig. S10. Interaction analysis of *At*SAR1c-GFP with endogenous *At*PHF1 and *At*PHT1;1/2/3.** Eleven-day-old wild-type (Col-0), *UBQ10:*GFP, and *UBQ10:At*SAR1c-GFP seedlings were subjected to Pi deprivation (0 µM $KH_2PO_4$, 7 days of starvation). Fifty-fold more root protein extract expressing *At*SAR1c-GFP was used as input for GFP-Trap co-immunoprecipitation relative to that expressing GFP. Anti-S11 antibody was used to detect GFP fusion proteins. Representative results are shown from three independent experiments.

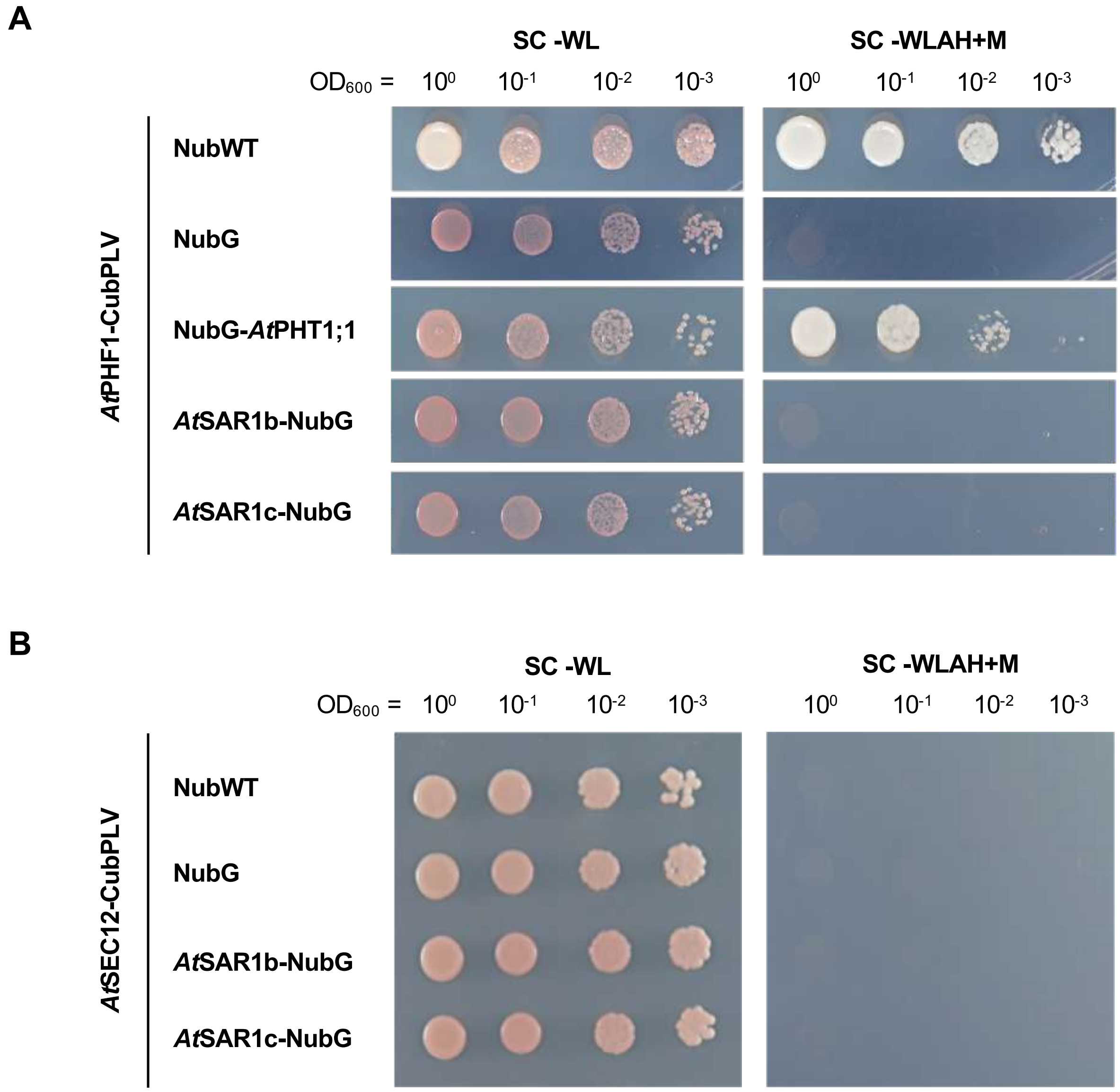


**Supplementary Fig. S11. Interaction analysis of *At*PHF1 with *At*SAR1 in the yeast split-ubiquitin system.** (**A**) Co-expression of *At*PHF1-CubPLV with NubG-*At*PHT1;1, *At*SAR1b-NubG or *At*SAR1c-NubG. (**B**) Co-expression of *At*SEC12-CubPLV with *At*SAR1b-NubG and *At*SAR1c-NubG. The co-expression of *At*PHF1-CubPLV or *At*SEC12-CubPLV with NubWT was used as a positive control, while the co-expression with NubG was used as a negative control. The CDS of *At*PHF1, *At*SEC12, *At*PHT1;1, *At*SAR1b, and *At*SAR1c were amplified by PCR, cloned into the pCR8/GW/TOPO, verified by sequencing, and recombined into destination vectors via LR reactions. *At*PHF1 and *At*SEC12 were cloned into synthesized MET17-PLVCub_GW to generate C-terminal Protein A–LexA–VP16–Cub fusions (*At*PHF1-CubPLV and *At*SEC12-CubPLV); *At*PHT1;1 was cloned into NX32_GW (ABRC plasmid CD3-1737) to produce NubG-*At*PHT1;1; *At*SAR1b and *At*SAR1c were cloned into XN22_GW (ABRC plasmid CD3-1735) to produce *At*SAR1b-NubG and *At*SAR1c-NubG. Yeast transformants were grown on synthetic medium lacking tryptophan and leucine (SC–WL) for growth detection or on synthetic medium lacking tryptophan, leucine, adenine, and histidine containing 0.5 µM methionine (SC–WLAH + M) for testing protein-protein interaction. Representative results are shown from two independent experiments.

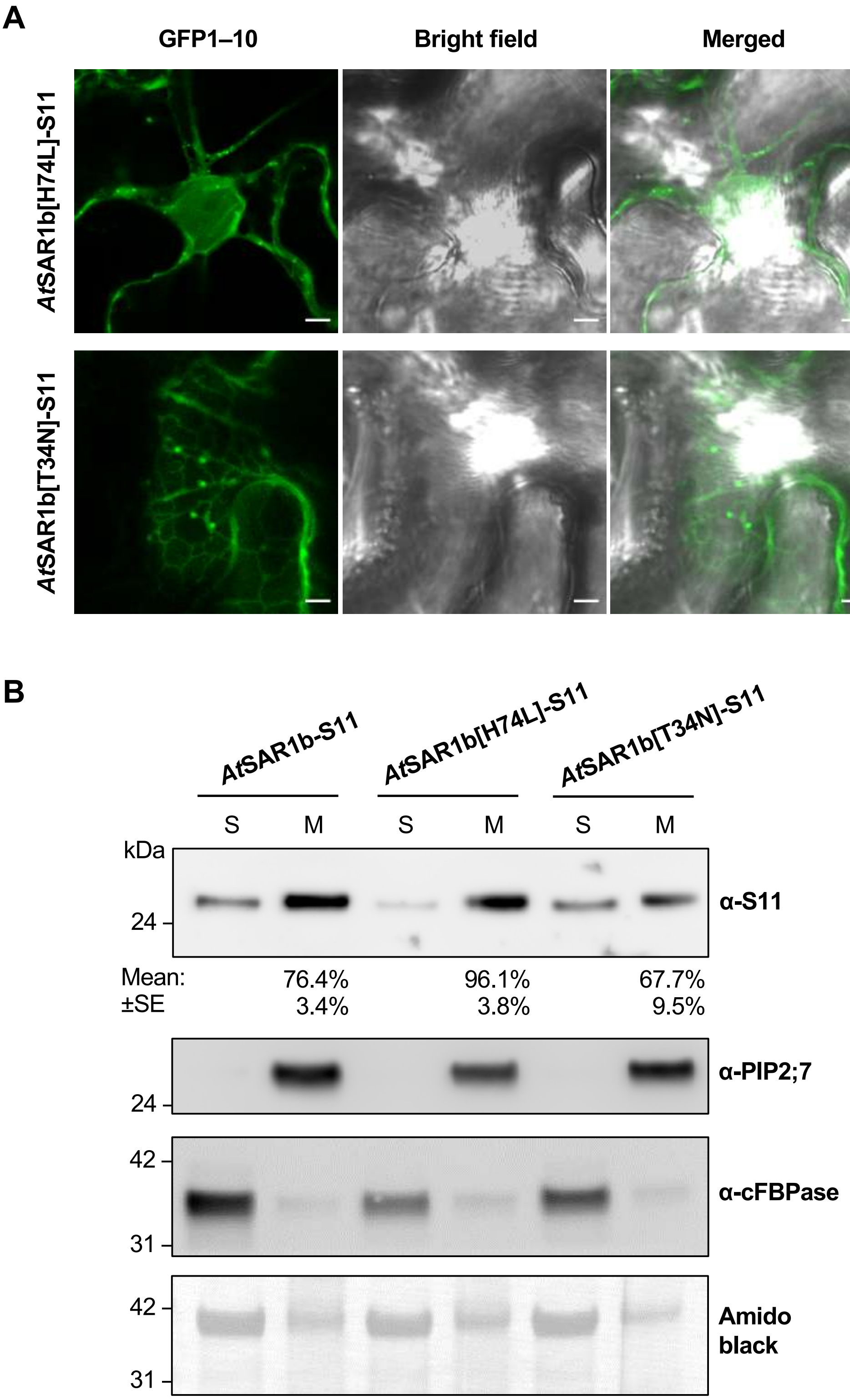


**Supplementary Fig. S12. Distribution of *At*SAR1b/[H74L]/[T34N] in the agro-infiltrated *N. benthamiana* leaves.** (**A**) The expression and distribution of the S11-tagged GTP-locked ([H74L]) and GDP-locked *At*SAR1b ([T34N]) at the peripheral layer of epidermal cells. Scale bars, 5 µm. Representative confocal images are shown from three independent experiments. (**B**) Protein expression of *At*SAR1b/[H74L]/[T34N] in the soluble (S) and microsomal (M) fractions isolated using the modified low-speed pellet (LSP) method . Anti-S11 antibody was used to detect S11 fusion proteins. The membrane association percentage (%) was calculated as M/(S+M) and is shown as mean ± standard error (SE) from two independent experiments. Aquaporin PIP2;7 and cytosolic fructose-1,6-bisphosphatase (cFBPase) served as microsomal and soluble controls, respectively. Amido black staining was used as a loading control.

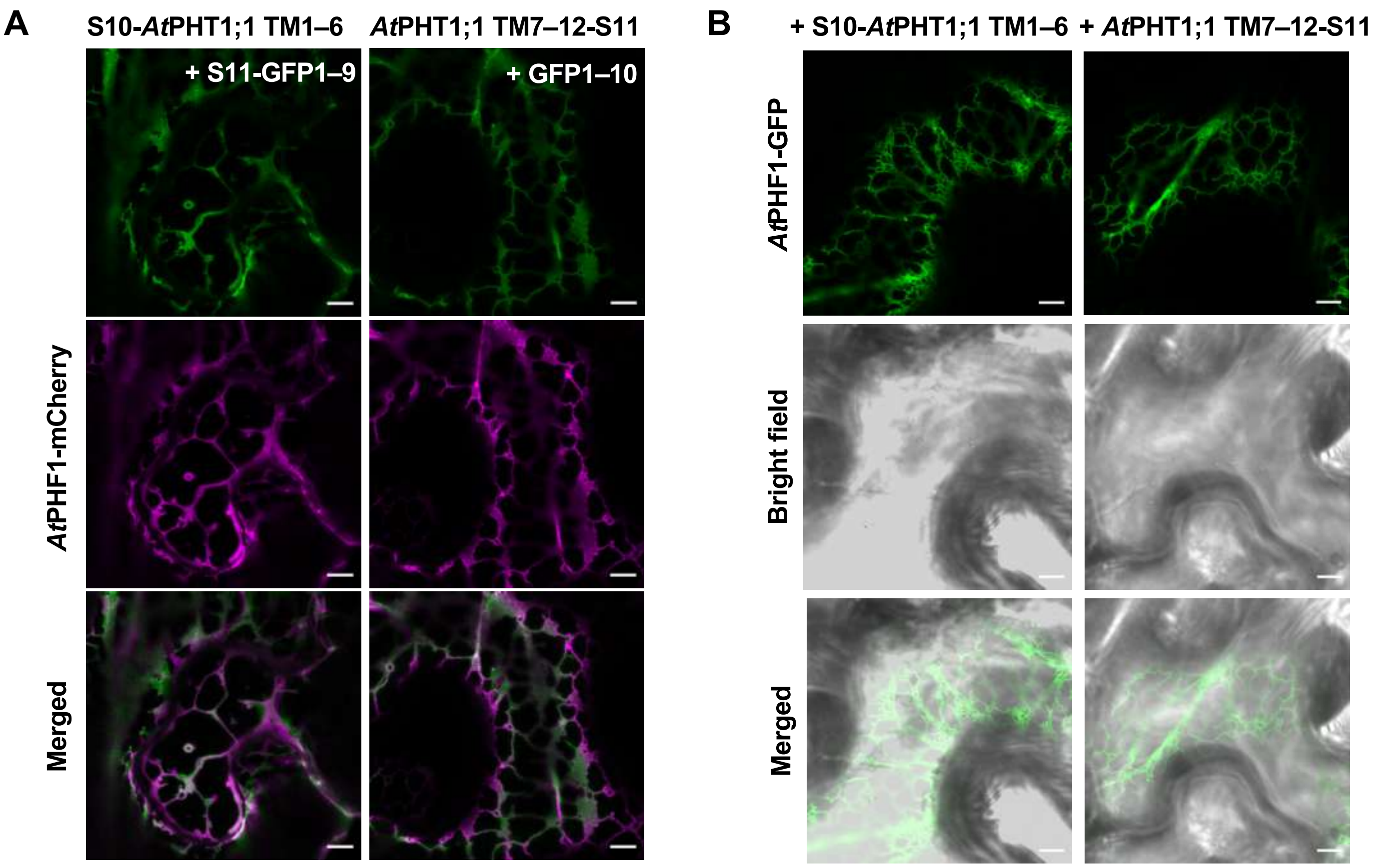


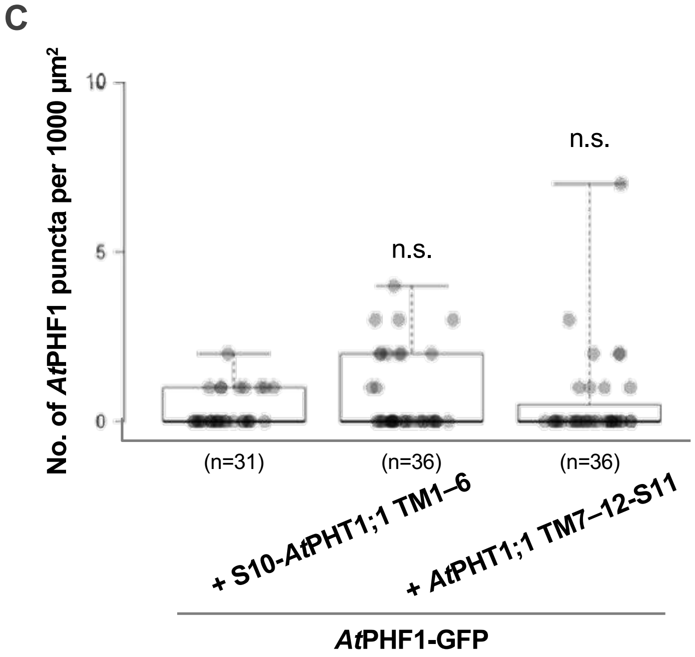


**Supplementary Fig. S13. ER export-incompetent *At*PHT1;1 overexpression cannot trigger partial distribution of *At*PHF1 to punctate structures in agro-infiltrated *N. benthamiana* leaves.** (**A**) Expression and distribution of S10-*At*PHT1;1 TM1–6 and *At*PHT1;1 TM7–12-S11 upon co-expression with *At*PHF1-mCherry in agro-infiltrated tobacco epidermal cells. Scale bars, 5 μm. Representative confocal images taken at the peripheral layer of epidermal cells are shown from three independent experiments. (**B**) Distribution of *At*PHF1-GFP in the presence of S10-*At*PHT1;1 TM1–6 or *At*PHT1;1 TM7–12-S11 co-expression. Scale bars, 5 μm. Representative confocal images are shown from three independent experiments. (**C**) Quantification of *At*PHF1-GFP-labeled punctate structures in (**B**). The data for *At*PHF1-GFP expressed alone are the same as those shown in Fig. 1B. Box plots display medians (center lines), interquartile ranges (boxes), and data ranges (whiskers), along with individual data points (dots). The number of regions of interest used for quantification is shown in parentheses. Dunnett's test for multiple comparisons against *At*PHF1-GFP expressed alone. Not significant, n.s..

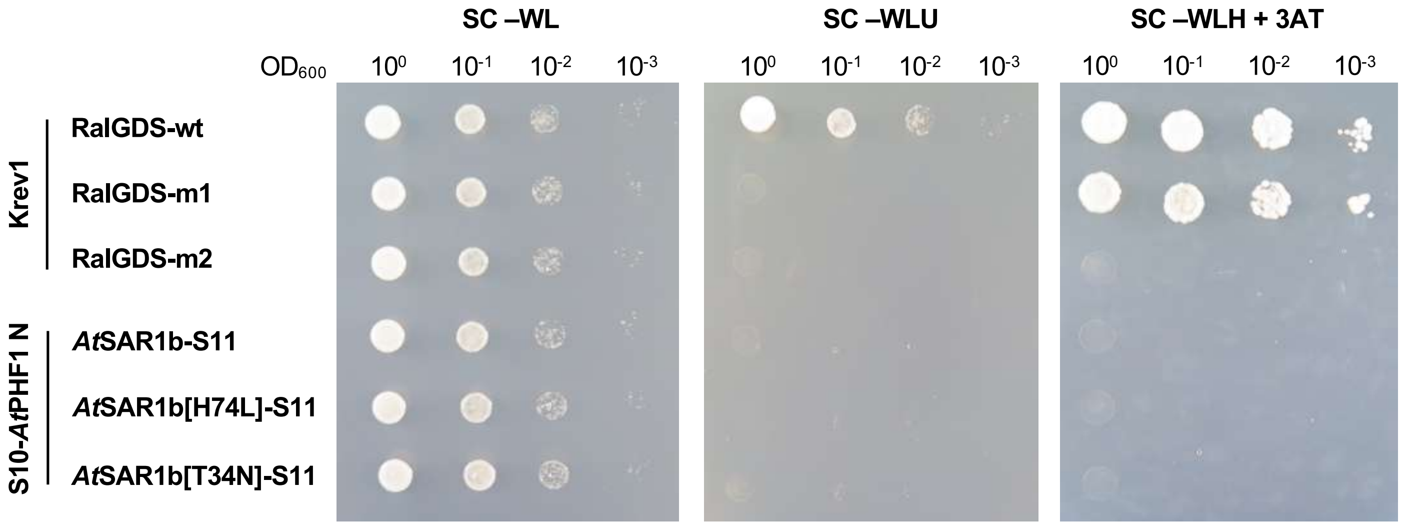


**Supplementary Fig. S14. Interaction analysis of the N-terminal cytosolic domain of *At*PHF1 with *At*SAR1b/[H74L]/[T34N] in the yeast two-hybrid assay.** GAL4[BD]-S10-*At*PHF1 N was co-expressed with GAL4[AD]-*At*SAR1b-S11, GAL4[AD]-*At*SAR1b[H74L]-S11, or GAL4[AD]-*At*SAR1b[T34N]-S11. The co-expression of Krev1 with RalGDS-wt, RalGDS-m1, and RalGDS-m2 were strong positive, weak positive, and negative controls, respectively. The CDS of *At*SAR1b-S11, *At*SAR1b[H74L]-S11, and *At*SAR1b[T34N]-S11 were subcloned into pCR8/GW/TOPO and recombined into pDEST22 via LR reaction to generate GAL4[AD]-*At*SAR1b/[H74L]/[T34N]-S11; the CDS of S10-*At*PHF1 N was subcloned into pCR8/GW/TOPO and recombined into pDEST32 via LR reaction to generate GAL4[BD]-S10-*At*PHF1 N. Yeast transformants were grown on synthetic medium lacking tryptophan and leucine (SC–WL) for growth detection, and on medium lacking tryptophan, leucine, and uracil (SC–WLU) or lacking tryptophan, leucine, and histidine with 10 mM 3-Amino-1,2,4-triazole (SC–WLH + 3AT) to assess protein-protein interactions. Representative results are shown from two independent experiments.

**Supplementary Table S1. Oligonucleotides used for cloning.**

| Gene/AGI No. | Primer name | Sequence (5' to 3') |
|---|---|---|
| *At*PHF1/AT3G52190 | AscI_AtPHF1.for1 | ggcgcgccaATGGAGATTGAAGAAGCGAGTCG |
| | AtPHF1 noSTOP_XhoI.rev | ctcgagTAGGTCCAAGTTCCACCTACTATG |
| | AtPHF1_PacI.rev | ttaattaaAGGTCCAAGTTCCACCTACTATGAT |
| *At*PHF1 truncated variants | AtPHF1(aa335)_XhoI. rev | CTCGAGCTCTTTCCACTCCTTTGGTACAGT |
| | AtPHF1(aa367)_XhoI.rev | ctcgagTGGTAACTTCCAAAACGAATCT |
| | AscI_AtPHF1(aa313).for | ggcgcgccaATGCTGACAACTTCTAGCGAAT |
| *At*SEC12/AT2G01470 | AscI_AtSEC12.for | ggcgcgccaATGGCGAATCAGAGTACAGAGA |
| | AtSEC12 STOP_PacI.rev | ttaattaaCTAAGGTATGATACCCTTTGCCT |
| | AtSEC12_PacI.rev | ttaattaaAGGTATGATACCCTTTGCCT |
| *At*PHT1;1/AT5G43350 | XbaI_AtPHT1;1.for | tctagaATGGCCGAACAACAACTAG |
| | AtPHT1;1 noSTOP_SpeI.rev | actagtTTTCTCGTCATGGCTAACCTC |
| | Infusion.F1 | caaatctaggggatccATGGCCGAACAACAACTAGGAGTGC |
| | Infusion.R1 | tttagctagactcgagTTATTTCTCGTCATGGCTAACCTCAGCC |
| *At*PHT1;1 truncated variants | AscI_AtPHT1;1 CDS.for | ggcgcgccaATGGCCGAACAACAACTAG |
| | AtPHT1;1(aa273)STOP_XhoI.rev | ctcgagtTTAATCCTCCACCCTTTCCT |
| | AscI_Start AtPHT1;1(aa273).for | ggcgcgccaATGGATGACGTCAAAGACCCC |
| | AtPHT1;1 CDS noSTOP_XhoI.rev | ctcgagTTTCTCGTCATGGCTAACCTC |
| *At*STP1/AT1G11260 | AscI_AtSTP1.for | aaggcgcgccaATGCCTGCCGGTGGATTCGTC |
| | AtSTP1 noSTOP_XhoI.rev | ctcgagAACATGCTTCGTTCCAGCTTGGT |
| *At*SAR1b/AT1G56330 | AscI_AtSAR1b.for | aaggcgcgccatgttcttgttcgattg |
| | AtSAR1b noSTOP_XhoI.rev | ctcgaggttgatgtactgagagagccatttgaatc |
| | Infusion_SmaI.F1 | gagggtaccgctcccgggatcccgctgctgaacgctaaa |
| | Infusion_XhoI.R1 | gactcgagcttttcggcagaccgca |
| | Infusion_XhoI.F2 | cgaaaagctcgagtctagtGGTGGAGGGTC |
| | Infusion_EcoRI.R2 | CAGCCGATCTGAATTCCCCGATCTA |
| *At*SAR1b[T34N] | Infusion 15bp overlapping_XbaI.F1 | tctgattaactctagactcgactctagcca |
| | Infusion 20bp overlapping_T34N.R1 | aagtaatgtgtttttgccagcattatcgagaccga |
| | Infusion 20bp overlapping_aa35.F2 | ctggcaaaaacacattacttcacatgctcaaagacga |
| | Infusion 20bp overlapping_XhoI.R2 | TCCACCactagactcgaggtt |
| *At*SAR1c/AT4G02080 | AscI_AtSAR1c.for | aaggcgcgccatgttcatgatcgattg |
| | AtSAR1c noSTOP_XhoI.rev | ctcgagcttgatgtattgagaaacccattt |
| | EcoRI_AtSAR1c.for | aagaattcatgttcatgatcgattggttc |
| | AtSAR1c (noSTOP)_EcoRI.rev | gaattccttgatgtattgagaaacccattt |
| *At*SEC24a/AT3G07100 | AscI_AtSEC24a.for | ggcgcgccatgggtacggagaatca |
| | AtSEC24a noSTOP_XhoI.rev | ctcgaggttttgttgaacttggcggtg |
| *At*SEC13a/AT3G01340 | AscI_AtSEC13a.for | aaggcgcgccatgccaggtcagaagat |
| | AtSEC13a noSTOP_XhoI.rev | ctcgagaggctcaacagcagtaacttgtt |
| *At*SEC16a/AT5G47480 | AscI_AtSEC16a.for | ggcgcgccatggcttcgactgctga |
| | AtSEC16a noSTOP_SalI.rev | gtcgaccagttcaacttcctgaagctcctctcc |

| | | |
|---|---|---|
| GFP/mCherry fusion | UBQ10 pro_HindIII.for | aagcttcgacgagtcagtaataaa |
| | UBQ10 pro_XbaI.rev | ctcgagtctagagttaatcagaaaaactcagatt |
| | SmaI_mCherry.for1 | cccgggATGGTGAGCAAGGGCGAGGA |
| | mCherry_SacI.rev1 | gagctcCTACTTGTACAGCTCGTCCATG |
| | GFP1-9 V16I_ApaI.for | aagggcccATGAGAAAAGGAGAAGAACTTT |
| | GFP196_SacI.rev | gagctcttaTGGTAAAAGGACAGGGCCATCG |
